# Allosteric modulation of TIA-1 phase separation by double serine phosphorylation

**DOI:** 10.1101/2025.08.21.671602

**Authors:** Alejandro Velázquez-Cruz, Laura Corrales-Guerrero, Ana B. Uceda-Mayo, Emanuela Tumini, Sofía M. García-Mauriño, Saboora Waris, Rafael L. Giner-Arroyo, Fionna E. Loughlin, Rafael Fernández-Chacón, Jacqueline A. Wilce, Miguel A. De la Rosa, Irene Díaz-Moreno

**Author notes:** Institute of Plant Biochemistry and Photosynthesis, — Centro de Investigaciones Científicas Isla de la Cartuja (cicCartuja), Universidad de Sevilla – Consejo Superior de Investigaciones Científicas (CSIC), Seville, Spain. Department of Chemistry, University of the Balearic Islands – Health Research Institute of the Balearic Islands (IdISBa) – Institut Universitari d’Investigació en Ciències de la Salut (IUNICS), Palma de Mallorca, Spain (Ana B. Uceda-Mayo).

## Abstract

In response to diverse harmful stimuli, eukaryotic cells generate cytoplasmic stress granules (SGs), mainly composed of mRNAs and RNA-binding proteins (RBPs). RBPs are fine-tuned by a diverse array of post-translational modifications (PTMs), with important consequences for the assembly, dynamics and clearance of SGs. One of the best characterized SG nucleators is the RBP T-cell intracellular antigen 1 (TIA-1), although knowledge about the structural and functional impact of its identified PTMs is very limited. TIA-1 is organized into three RNA-recognition motifs (RRMs) and a C-terminal prion-related domain (PRD) that drives its phase separation from the cytosol. Here, we analyzed the effect of TIA-1 double phosphorylation in RRM3, at serines 198 and 199. Microscopic observations revealed an increased propensity of the phosphomimetic TIA-1 S198/199E to undergo liquid-liquid phase separation (LLPS) and self-assemble into SGs independently of stress stimuli. Our computational simulations, supported by NMR data, have suggested that such phosphorylations promote the formation of a β-hairpin motif at the beginning of the PRD. Moreover, the ALS-associated mutation V283M in TIA-1 was predicted to lead to the formation of an aberrant structure in the β-hairpin region, highlighting the fine balance between physiological and pathogenic TIA-1 phase transition, and the importance of a better understanding of the molecular mechanisms underlying the liquid demixing of this RBP.

## Introduction

Stressed cells undergo a global depletion of protein synthesis and focus on the expression of pro-survival genes^1^. Under such circumstances, a conserved event among eukaryotes is the rapid accumulation of mRNA and their cognate RNA-binding proteins (RBPs) to form the so-called stress granules (SGs)^2,3^. SGs are cytoplasmic condensates of variable shape, structure and composition, instigated by diverse harmful stimuli (e.g., arsenite, heat shock and UV radiation) and whose typically ascribed role is the transitory storage of non-essential and untranslating mRNA pools, conceivably supporting the optimal management of cell energy resources and a prompt return to basal conditions upon stress removal^1,3–5^. Additionally, SGs have emerged as versatile regulatory hubs to cope with different adverse situations, modulating cellular stress responses by sequestering key proteins of signaling pathways^6,7^.

The formation of these membrane-less organelles is strongly correlated with polysome disassembly and release of mRNA-containing translation initiation complexes^8^, which serve as scaffolds for the recruitment of nucleating RBPs such as T-cell intracellular antigen-1 (TIA-1) and Ras GTPase-activating protein SH3 domain-binding protein 1 (G3BP1)^6,9,10^, commonly used SG markers in cell-based assays. Importantly, most SG nucleators possess intrinsically disordered regions (IDRs) that engage in multivalent homotypic and/or heterotypic protein-protein interactions, contributing prominently to the liquid-liquid phase separation (LLPS) of messenger ribonucleoprotein complexes^5,6^.

The LLPS of certain proteins can favor their aggregation into amyloids^11^. Indeed, the dysregulation of SGs dynamics leads to the development of a great variety of diseases^5,10^. Pathological SGs are more frequently related to neurodegenerative disorders, such as amyotrophic lateral sclerosis (ALS) and frontotemporal dementia (FTD), characterized by an imbalance between SGs formation and dismantling, as well as the liquid-to-solid maturation of SGs^12–14^. For example, several mutations have been linked to the higher propensity of TIA-1 to undergo LLPS and fibrillization in ALS/FTD patients, giving rise to low-dynamic and persistent SGs^15^.

SGs are extensively controlled by post-translational modifications (PTMs) on their integral proteins, decisively influencing the generation, dynamics and clearance of these biomolecular condensates^5,16^. There is abundant literature on PTMs regulating the stability, aggregation state (dispersed or phase-separated), interactions and subcellular localization of SG-associated RBPs^17,18^. For instance, specific serine phosphorylations of G3BP1 and lysine acetylations of Fused in Sarcoma (FUS) protein have been shown to impair their RNA-induced condensation^19–21^. In addition, arginine methylation in the RGG-rich regions of the aforementioned SG nucleators was also demonstrated to severely reduce their LLPS capability^22–24^. In contrast, serine phosphorylations in the C-terminal intrinsically disordered domain of both FUS and TIA-1 related protein-2 (TIAR-2) increase their LLPS in living cells^25,26^. Likewise, lysine acetylation in the nuclear localization signal (NLS) of FUS leads to its cytoplasmic accumulation and enhanced liquid demixing^19^. Interestingly, hypomethylated and NLS-acetylated FUS is characteristic of FTD^19,23,24^.

The well-studied RBP TIA-1 generally recognizes A/U- or C/U-rich sequences in untranslated regions of precursor and mature mRNA targets, modulating alternative splicing in the nucleus and translation in the cytoplasm^17,27–30^. Furthermore, TIA-1 also exhibits high affinity for T-rich single-stranded (ss)DNA^28^, hinting at the participation of this RBP in the functional coupling between transcription and splicing^31^. TIA-1 structure consists of three RNA-recognition motifs (RRMs) and a C-terminal prion-related domain (PRD) that drives its phase separation from the cytosol (**Figure 1A**)^32^. Nevertheless, the RNA-binding ability of TIA-1 is indispensable for its recruitment into SGs^8^, and it has been documented that nucleic acids harboring multiple binding sites for TIA-1 can induce its LLPS and further aggregation into amyloid-like fibrils^33^. Interestingly, Zn^2+^ and RGG motifs have recently been revealed as co-factors that not only promote the phase separation of TIA-1 but also inhibit its transition into fibrillar aggregates, representing a protective mechanism against fibrillization in the cell^34,35^.

**Figure 1.**
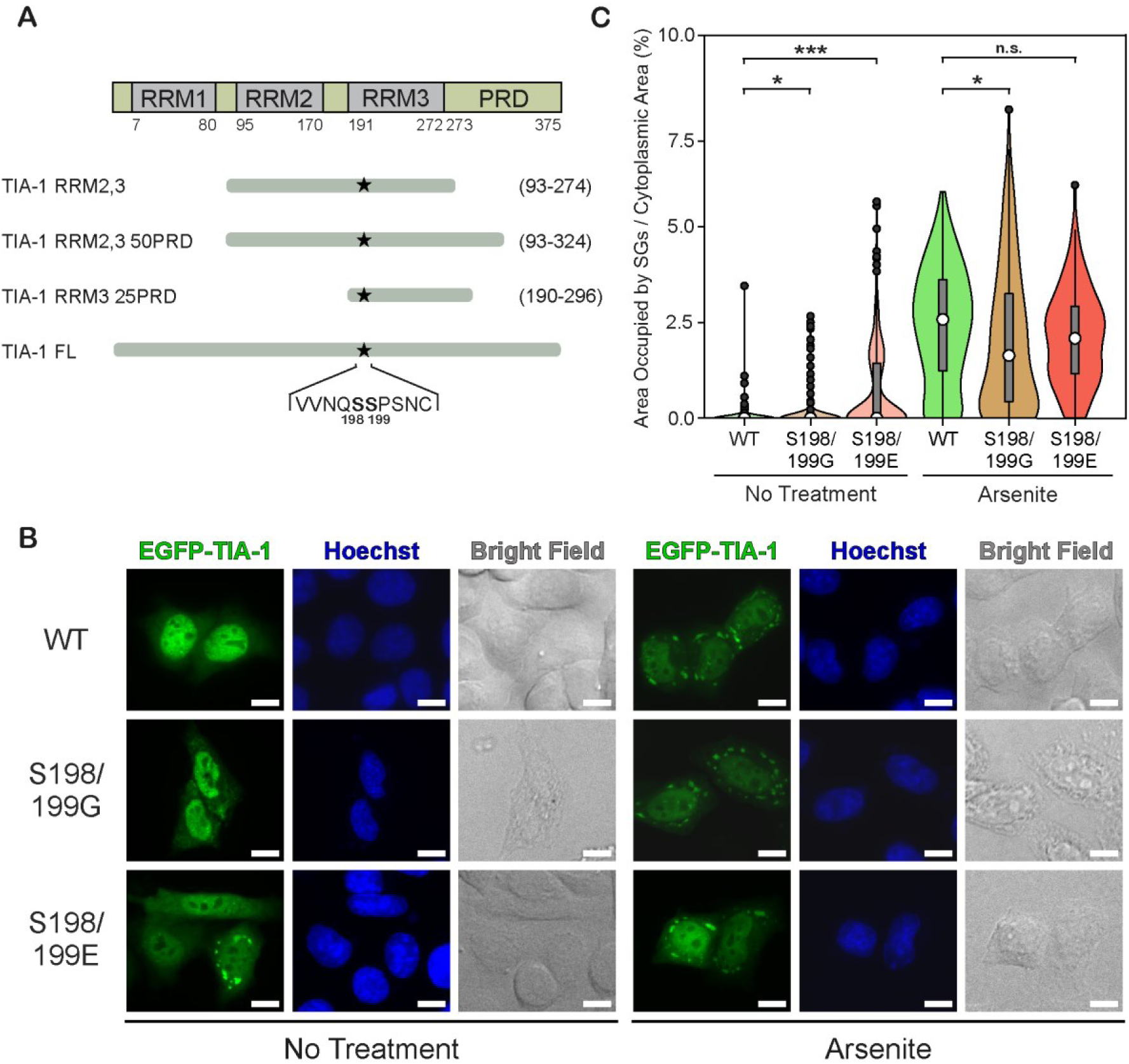
Phosphomimetic TIA-1 S198/199E spontaneously assembles into SGs. **A)** TIA-1 isoform b domain organization and constructs used in the present study, highlighting the location of the phosphorylatable serines 198 and 199. **B)** Representative fluorescence microscopy images of HeLa cells transfected with different EGFP-TIA-1 constructs (green fluorescence) under homeostatic and oxidative stress conditions. Nuclei were stained with Hoechst (blue fluorescence). Scale bars are 10 μm. **C)** The area of the cytoplasm occupied by SGs for each EGFP-TIA-1 construct assayed, expressed by non-treated or arsenite-treated cells, is shown as a violin plot, with the white dot indicating the median; the black dots, the outliers; the box, the interquartile range; and the whiskers extending to the lowest and highest value within 1.5 times, the interquartile range from the hinges. In addition, rotated kernel density plots are given for each dataset to show their distribution. N = 300 cells for basal conditions and 150 cells for stress conditions. The significance of differences was tested using the Student’s t test. n.s., not significant; \**p* < 0.05; \*\*\**p* < 0.001.

As explained before, the context-dependent condensation or aggregation of RBPs is highly regulated by PTMs, but limited information exists in this regard for TIA-1 (PhosphoSitePlus database^36^). Here we have investigated the possible impact on TIA-1 SG-nucleating activity of its phosphorylation at serines 198 and 199, first reported in a proteomic study^37^. In order to characterize the structural and functional properties of S198/199-phosphorylated (pS198/199) TIA-1, we resorted to a multidisciplinary approach that combined cell biology, biophysics and computational methods. Our results uncovered a potential allosteric pathway governing TIA-1 LLPS, by which the double serine phosphorylation of RRM3 would trigger the assembly of a β-hairpin motif at the beginning of the PRD that facilitates the liquid demixing of TIA-1 and, therefore, SGs formation.

## Results

### Phosphomimetic TIA-1 S198/199E displays enhanced phase-transition propensity

To explore the possible physiological significance of TIA-1 S198/199 phosphorylation, HeLa cells expressing EGFP-tagged TIA-1 full-length WT, S198/199E (phosphomimetic) and S198/199G (non-phophorylatable) constructs were examined by fluorescence microscopy (**Figure 1**). Under basal conditions, an unusually high occurrence of SGs in cells expressing the phosphomimetic mutant of TIA-1 was observed, in contrast to the practical absence of these biomolecular condensates in cells transfected with the plasmids for WT and non-phosphorylatable TIA-1 variants (**Figure 1B**). In order to corroborate these observations, a systematic quantification was accomplished for each TIA-1 construct, consisting of the analysis of the cytoplasmic area occupied by SGs in a representative subset of cells. In parallel, expression levels were also estimated based on total cell fluorescence (**Supplementary Figure S1A**), showing a similar range of concentrations for all TIA-1 variants under the same experimental conditions. The quantification results demonstrated that phosphomimetic TIA-1 S198/199E has an abnormal tendency to form SGs in the absence of stress stimuli (**Figure 1B-C**). However, under arsenite-induced oxidative stress, the presence of SGs was rather uniform among all constructs assayed (**Figure 1B-C**), suggesting that S198/199 phosphorylation is not essential for TIA-1 assembly into SGs.

To further characterize the impact of S198/199 phosphorylation on the LLPS propensity of TIA-1, turbidity measurements and droplet formation assays were performed using purified TIA-1 full-length WT and S198/199E constructs. Analogously, absorbance readings at 385 nm showed an elevated intrinsic aggregation capacity of the phosphomimetic TIA-1 mutant compared to WT, both in the absence and presence of the multisite TC5 ssDNA (that has previously been shown to induce LLPS on TIA-1^33^) (**Figure 2A**). Complementary droplet formation assays were carried out using DIC microscopy. Once again, the role of phosphomimetic mutations in the promotion of free TIA-1 condensation was clearly revealed (**Figure 2B-C**). Furthermore, addition of TC5 ssDNA to TIA-1 samples produced a drastic increase in the number of droplets, but the S198/199E variant still showed a higher propensity for droplet formation than the WT protein (**Figure 2B-C**). It can therefore be concluded that the mechanism favoring TIA-1 S198/199E phase transition does not require nucleic acid binding. Taken together, experimental evidence points to S198/199 phosphorylation as a potential biochemical switch contributing to TIA-1 LLPS. This is in agreement with that reported for TIAR-2, whose PRD serines are not only phosphorylated *in vivo* but also promote the capacity of TIAR-2 to undergo LLPS^26^.

**Figure 2.**
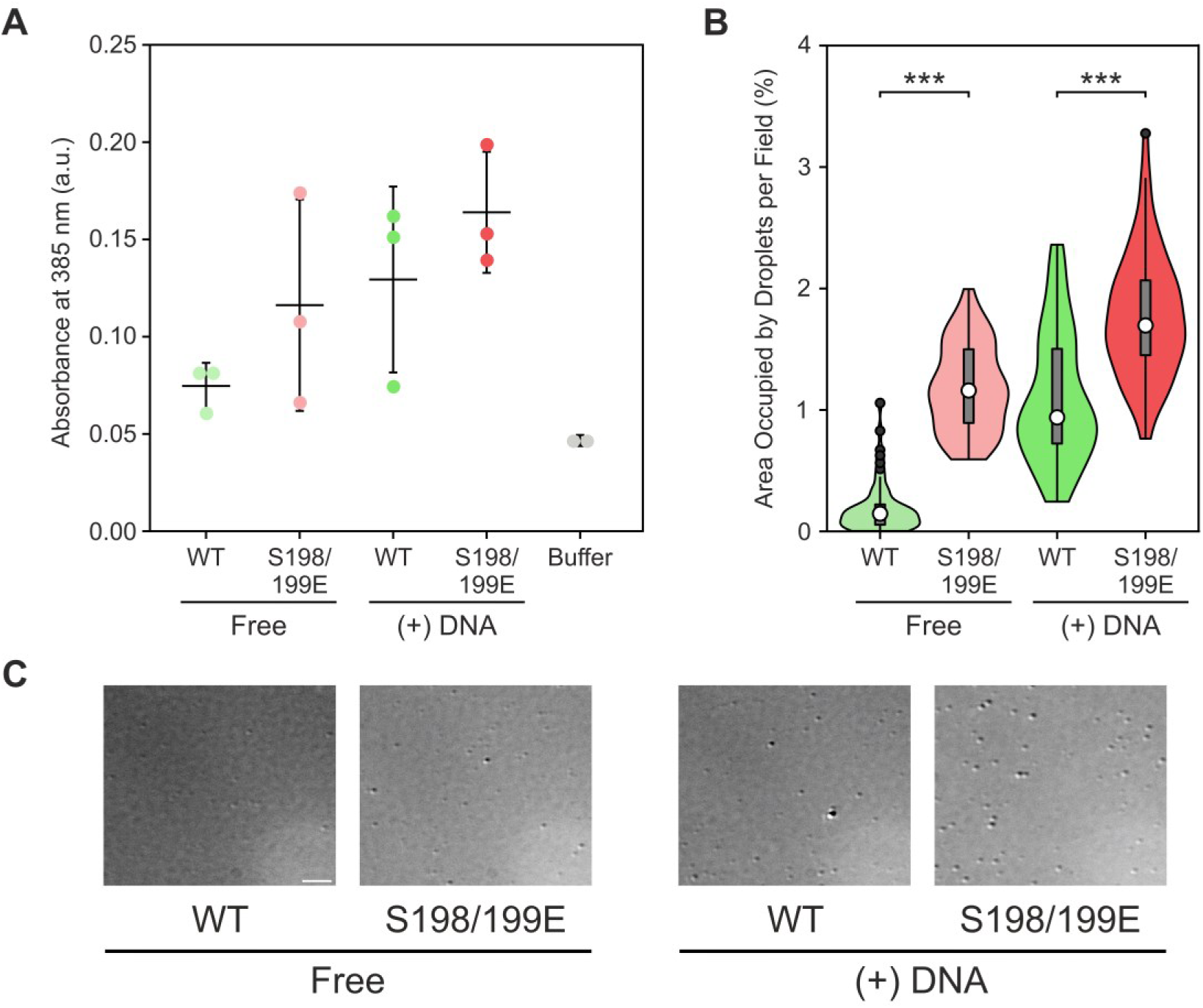
The increased LLPS propensity of TIA-1 S198/199E does not rely on the association with nucleic acids. **A)** Turbidity assay comparing the aggregation capacity of TIA-1 full-length WT and S198/199E in the presence and absence of the TC5 ssDNA oligo. Dots represent different replicates of each sample, with horizontal lines and error bars indicating the average and the standard deviation (S.D.), respectively. **B)** Droplet formation assay with the same protein constructs and conditions as in (**A**) showing the area occupied by droplets per field in a violin plot, with the white dot indicating the median; the black dots, the outliers; the box, the interquartile range; and the whiskers extending to the lowest and highest value within 1.5 times, the interquartile range from the hinges. In addition, rotated kernel density plots are given for each dataset to show their distribution. N = 60 fields from 3 replicates. The significance of differences was tested using the Student’s t test. \*\*\**p* < 0.001. **C)** Representative bright-field microscopy images of the samples tested in (**B**). Scale bar is 5 μm.

### Phosphomimetic double mutation S198/199E reshapes TIA-1 intramolecular interactions between RRM3 and PRD

Since TIA-1 assembly into SGs is mainly driven by PRD self-association^32^, phosphomimetic glutamate mutations and, by extension, phosphorylations at S198/199 could be somehow facilitating the homotypic multimerization of the PRD. In order to study the possible conformational changes in TIA-1 derived from S198/199 phosphorylation or substitution for glutamate, we employed a hybrid biophysical and *in silico* approach. Serines under examination are located between the α_0_-helix and β_1_-strand of TIA-1 RRM3, partially burying the nearby residue W272, which is at the C-terminal of the same domain^29,30,38^. This configuration, together with the intrinsic fluorescence properties of tryptophan, makes W272 an ideal endogenous probe to register any structural rearrangement around the phosphomimetic mutations. Therefore, these unique features of TIA-1 were exploited through fluorescence spectroscopy measurements, as previously reported^38,39^. But first, three out of four tryptophans — W147, W160 and W170 — in TIA-1 RRM2,3 were replaced by phenylalanine, so that only W272 remained in the constructs tested. Of note, these new TIA-1 species were found to retain the native folding of the tandem domains by CD measurements (**Supplementary Figure S2A-B**). Although the fluorescence spectra of both TIA-1 triple tryptophan-to-phenylalanine mutants, S198/199 and E198/199, hardly showed differences in the maximum emission wavelength, the phosphomimetic mutations produced a large increase in the fluorescence intensity of W272 (**Figure 3A**). The observed small red-shift (∼2 nm) resulting from S198/199E substitutions denotes a similar degree of solvent exposure for W272 in both constructs used^40^. However, the remarkable variation in fluorescence quantum yield could be indicative of critical changes in TIA-1 RRM3 structure due to serine-to-glutamate mutations^41,42^. Subsequent monitoring of TIA-1 RRM2,3 W147/160/170F denaturation by urea titration showed quenching of W272 fluorescence in the native state of this construct (**Supplementary Figure S2C**). Such loss of energy in the non-phosphomimetic version of TIA-1 could be triggered by electron-transfer from the excited indole ring of W272 to any neighboring peptide bond^43,44^. The fact that S198/199E mutations eliminate the quenching of W272 in the native folding of TIA-1, leading to an increase in the fluorescence intensity comparable to that of urea denaturation, supports the emergence of notable structural changes in RRM3 upon S198/199 phosphorylation.

**Figure 3.**
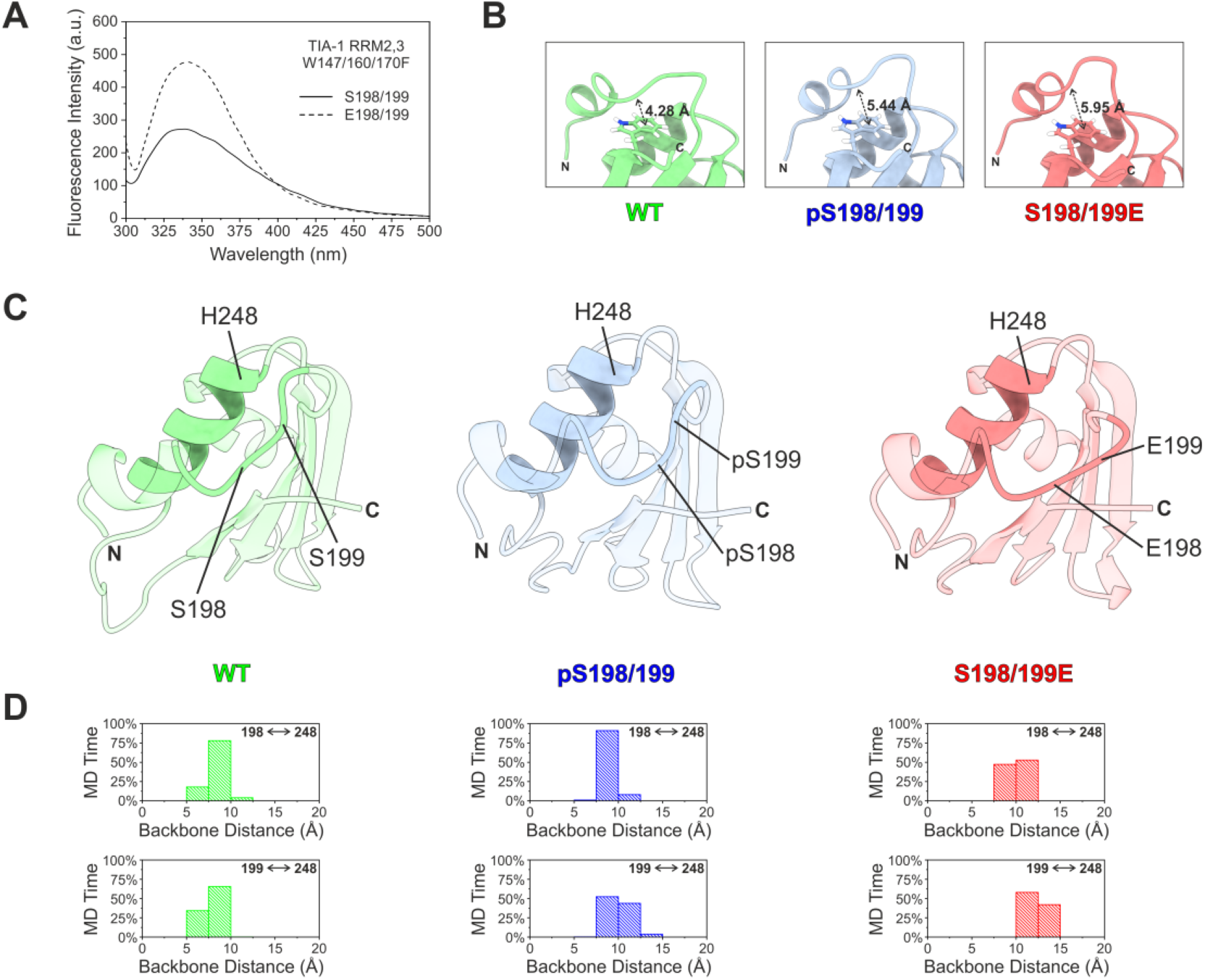
TIA-1 S198/199 phosphorylation and glutamate mutation give rise to structural changes in RRM3. **A)** Spectra of TIA-1 RRM2,3 W147/160/170F monitoring the fluorescence emission from W272, spatially close to residues 198/199 and used as a natural reporter of the structural rearrangements in its vicinity due to serine-to-glutamate mutations. **B-C)** Average structure of TIA-1 in the MD simulations, showing the distance between W272 and the loop connecting α_0_-helix (N-terminal) and β_1_-strand of RRM3 (**B**), as well as the opening of the latter region with respect to α_2_-helix (opaque ribbon stretches; for the sake of clarity, the rest of the domain is semi-transparent) (**C**). The distance between the centers-of-mass of the W272 benzene ring and the closest peptide bond is indicated in (**B**) for each TIA-1 model. **D)** Histogram plots of the distances between the centers-of-mass of the backbone atoms of residues 198/199 and H248 along the MD trajectories of TIA-1. H248, N-terminal residue of RRM3 α_2_-helix, was used as a reference to measure the relative displacement of S198/199 upon phosphorylation or phosphomimetic mutation.

To further explore the conformational impact of S198/199 phosphorylation at atomic detail, MD simulations were run using full-length models of TIA-1 WT, pS198/199 and S198/199E. The estimated average distance between W272 (indole ring) and residues 198/199 (backbone) along MD simulations was found to be increased due to serine phosphorylations or phosphomimetic mutations (**Figure 3B**), in good agreement with fluorimetry measurements. Moreover, comparison of the average TIA-1 RRM3 structure of each MD trajectory showed the S198/199-containing loop region bulging out of the domain and separating from adjacent α_2_-helix in the phosphorylated and phosphomimetic models (**Figure 3C-D**). Thus, computational analysis predicted that phosphorylations at S198/199 would produce a local rearrangement in TIA-1 RRM3 which, together with the expected shift in the surface electrostatic potential, could substantially interfere in both inter- and intramolecular interactions of this RBP.

In addition, it was noted that the S198/199-containing loop region approaches the initial stretch of the PRD during the MD simulations of phosphorylated and phosphomimetic models (**Figure 3C**), hence the establishment of new intramolecular interactions was analyzed. The main difference between TIA-1 species was found in the dynamics of S198 and K274, which bring their side chains closer to establish a strong polar interaction when the former residue is phosphorylated or mutated to glutamate (**Figure 4A-C**). Furthermore, attractive forces between pS198 or E198 and neighboring N202 (**Figure 4C**), often bound by a hydrogen bond (**Supplementary Figure S3A**), may facilitate the proper orientation of phosphoserine and glutamate side chains for interaction with K274 (**Figure 4A-B**). Interestingly, K274 can also associate with D278, especially in the phosphorylated TIA-1 model, thus acting as a ‘bridge’ between RRM3 and the PRD (**Figure 4A-C** and **Supplementary** Figure 3B). On the other hand, either pS199 or E199 would play a more secondary role in establishing inter-domain contacts (**Figure 4A,C**). Collectively, MD simulations anticipated that TIA-1 S198/199 phosphorylation and glutamate mutation would create new connections between RRM3 and the beginning of the PRD, perhaps as part of an unknown allosteric communication pathway controlling the arrangement of TIA-1 disordered domain.

**Figure 4.**
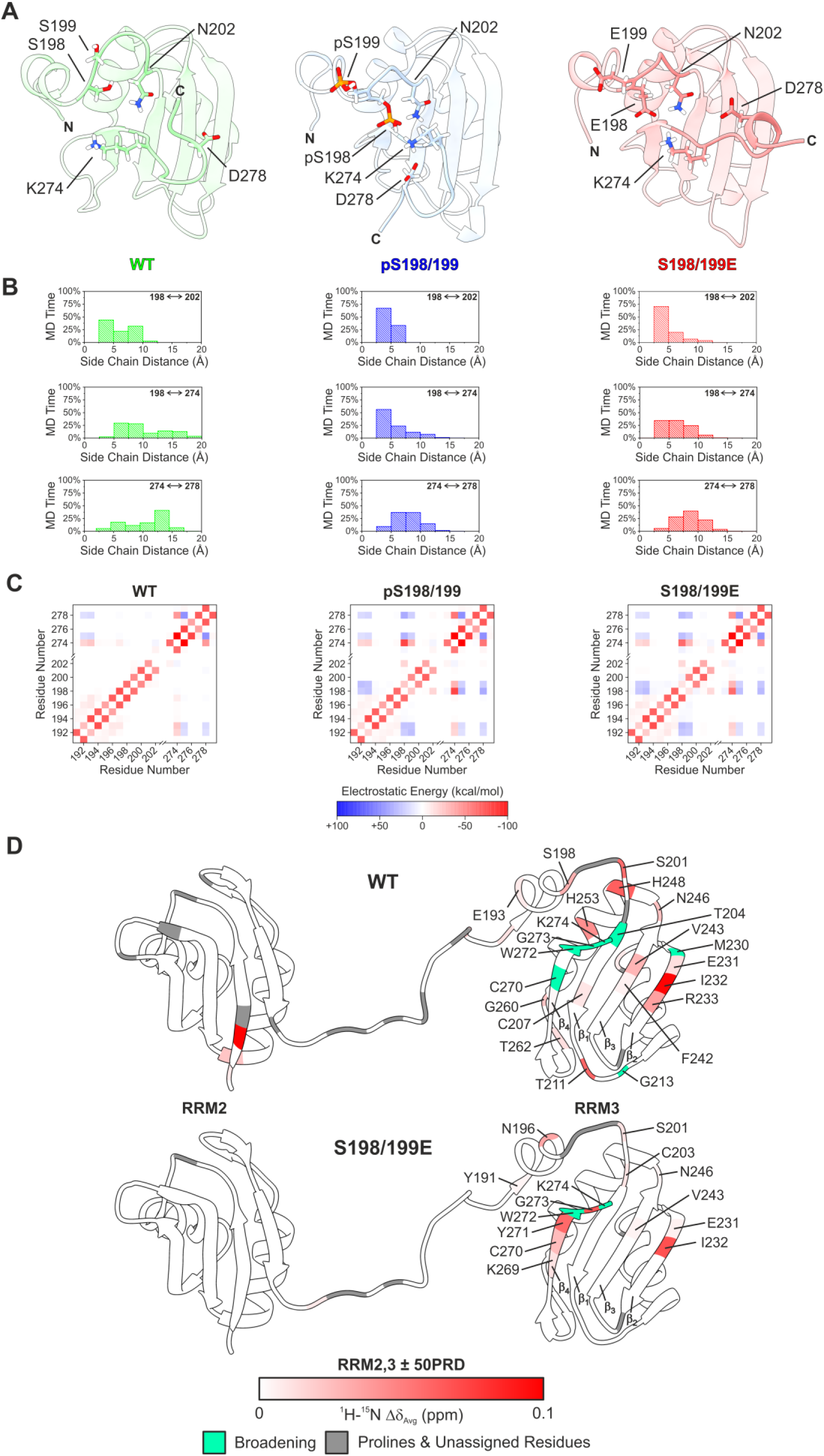
Remodeling of TIA-1 RRM3-PRD contacts by S198/199 phosphorylation and glutamate mutation. **A)** Representative ribbon structures of the most populated clusters from the MD simulations of TIA-1, showing RRM3 and the initial stretch of the PRD. Clustering of trajectory frames was based on RMSD (residues 191-297). **B)** Histogram plots of the distances between the centers-of-mass of the side-chain functional groups of residues 198, N202, K274 and D278, along the MD trajectories of TIA-1. **C)** Heatmaps showing the average electrostatic interaction energy between residues of the phosphorylation/mutation sites and the initial stretch of the PRD (residues between both regions have been omitted), generated from the MD simulations of TIA-1. The nature and magnitude of the electrostatic forces are represented by a diverging color palette, where attractive interactions appear in red and repulsive interactions in blue. **D)** Mapping of TIA-1 RRM2,3 residues (structure from ref. 29; PDB ID: 2MJN) perturbed by intramolecular contacts with the first half of the PRD (up to residue 324). Residues are colored according to their weighted-average chemical-shift perturbations (Δδ_Avg_) using a red gradient from 0 to 0.1 ppm. Residues with signals broadened beyond detection are indicated in green, whereas prolines and not assigned residues are in grey.

In order to investigate the effects of TIA-1 S198/199 phosphorylation on the presumed contacts between RRM3 and PRD, ^1^H-^15^N-HSQC NMR experiments were performed employing WT and phosphomimetic versions of ^15^N-labelled TIA-1 RRM2,3 and TIA-1 RRM2,3 50PRD (including the first 50 residues of the PRD) (**Figure 1A**). First, analysis of averaged chemical-shift perturbations (Δδ_Avg_) between the TIA-1 species devoid of PRD verified the impact of S198/199E mutations on their neighboring residues (**Supplementary Figure S4**), as expected from previous fluorometric and computational analyses. The comparison between spectra of TIA-1 constructs harboring or not ‘50PRD’ was used to assess the intramolecular interactions of the unstructured domain with RRM2,3. The negligible ^1^H-linewidth broadening (∼1-3 Hz on average) ascribable to 50PRD allowed us to exclude substantial TIA-1 self-association. Moreover, the vast majority of perturbed residues are located in RRM3 (**Figure 4D**), confirming the existence of intramolecular transient contacts between this domain and the PRD, whose initial stretch protrudes from the β-sheet face of RRM3. Mapping of Δδ_Avg_ onto TIA-1 RRM2,3 ribbon diagrams showed generally greater perturbations in WT than in S198/199E. Specifically, backbone amide signals of C207 (β_1_), E231 (β_2_), I232 (β_2_), R233 (β_2_), F242 (β_3_) and V243 (β_3_) exhibited higher Δδ_Avg_ values in WT when comparing the constructs with and without 50PRD. Furthermore, TIA-1 PRD sampling of RRM3 surface seemed to be restricted in the serine-to-glutamate mutant, where most affected resonances were clustered in the β_4_-strand (**Figure 4D** and **Supplementary Figure S5**). Overall, NMR data indicated that the TIA-1 disordered PRD establishes substantial transient intramolecular contacts with the β-sheet face of the folded RRM3, similarly as described for heterogeneous nuclear ribonucleoprotein A1 (hnRNPA1)^45^. These contacts are limited by the double mutation S198/199E, consistent with the aforementioned increased inter-domain connection that would generate a more rigid and ‘fixed’ N-terminal tract of the PRD for the mutant protein.

Altogether, the synergistic combination of computational and biophysical experiments uncovered a relatively subtle conformational rearrangement in TIA-1 RRM3 triggered by S198/199 phosphorylation, but with important consequences for the intramolecular interactions between this domain and the PRD module.

### TIA-1 S198/199 phosphorylation and glutamate mutation would promote the assembly of a β-hairpin motif in the PRD

In light of the growing implications of TIA-1 S198/199 phosphorylation and phosphomimetic mutation in the conformation and intramolecular interactions of the PRD, the MD simulations of TIA-1 full-length models were again analyzed, but this time with the focus on the unstructured domain. First, intramolecular contacts (< 6 Å) along the different trajectories were computed and displayed in heatmaps (**Supplementary Figure S6A**). In addition to corroborating the previously described RRM3-PRD contacts, these calculations yielded some intriguing results for the N-terminal tract of the disordered domain. In the MD simulations of both phosphorylated and phosphomimetic TIA-1, persistent contacts were detected approximately between residues 277-292, whose pattern matches two short β-strands (**Figure 5A**). In contrast, such contacts were completely absent in the trajectory of the WT model. Next, a complementary analysis of TIA-1 secondary structure across MD trajectories was performed (**Supplementary Figure S6B**). Residues P282, V283, N287 and Q288 were mostly assigned the β-strand category throughout the MD simulations of TIA-1 pS198/199 and S198/199E, whereas only V283 and, to a lesser extent, Q284 were considered as β-strand residues in the WT model (**Figure 5B**). Furthermore, all three versions of TIA-1 shared a short and highly unstable helix at the beginning of the PRD (**Figure 5B** and **Supplementary Figure S6C**). To better understand the structure predicted in the N-terminal quarter of the PRD upon S198/199 phosphorylation or mutation to glutamate, the distribution and lifetime of hydrogen bonds were thoroughly examined. Results of hydrogen bond analysis between residues 280-291 revealed a significantly higher occupancy in the phosphorylated and phosphomimetic models, especially in the first half of the stretch, where practically no hydrogen bonds were found in WT. More importantly, the hydrogen-bonding pattern of TIA-1 pS198/199 and S198/199E is consistent with that found in β-hairpin motifs^46,47^ (**Figure 5C**). The prevalent backbone hydrogen bond between V283 and N287, flanking three consecutive glutamines that form a turn, would presumably be indispensable for the assembly and stability of this β-hairpin^48–50^. Additional backbone hydrogen bonds such as I280-G290 and N281-I289, as well as side-chain hydrogen bonds (proton donor and/or acceptor from side chains) such as Q284-N287, Y291-N281 and G290-N281, would contribute, in varying degrees, to the consolidation of this structural motif (**Figure 5C-D**). Furthermore, the β-hairpin also forms hydrogen bonds with its adjacent helix (**Figure 5D** and **Supplementary** Figure 6D), thus configuring a network of electrostatic interactions that connects the N-terminal quarter of the PRD with pS198 or E198. Together with hydrogen bonding, interactions between hydrophobic residues must play a substantial role in the folding pathway of the β-hairpin^50^.

**Figure 5.**
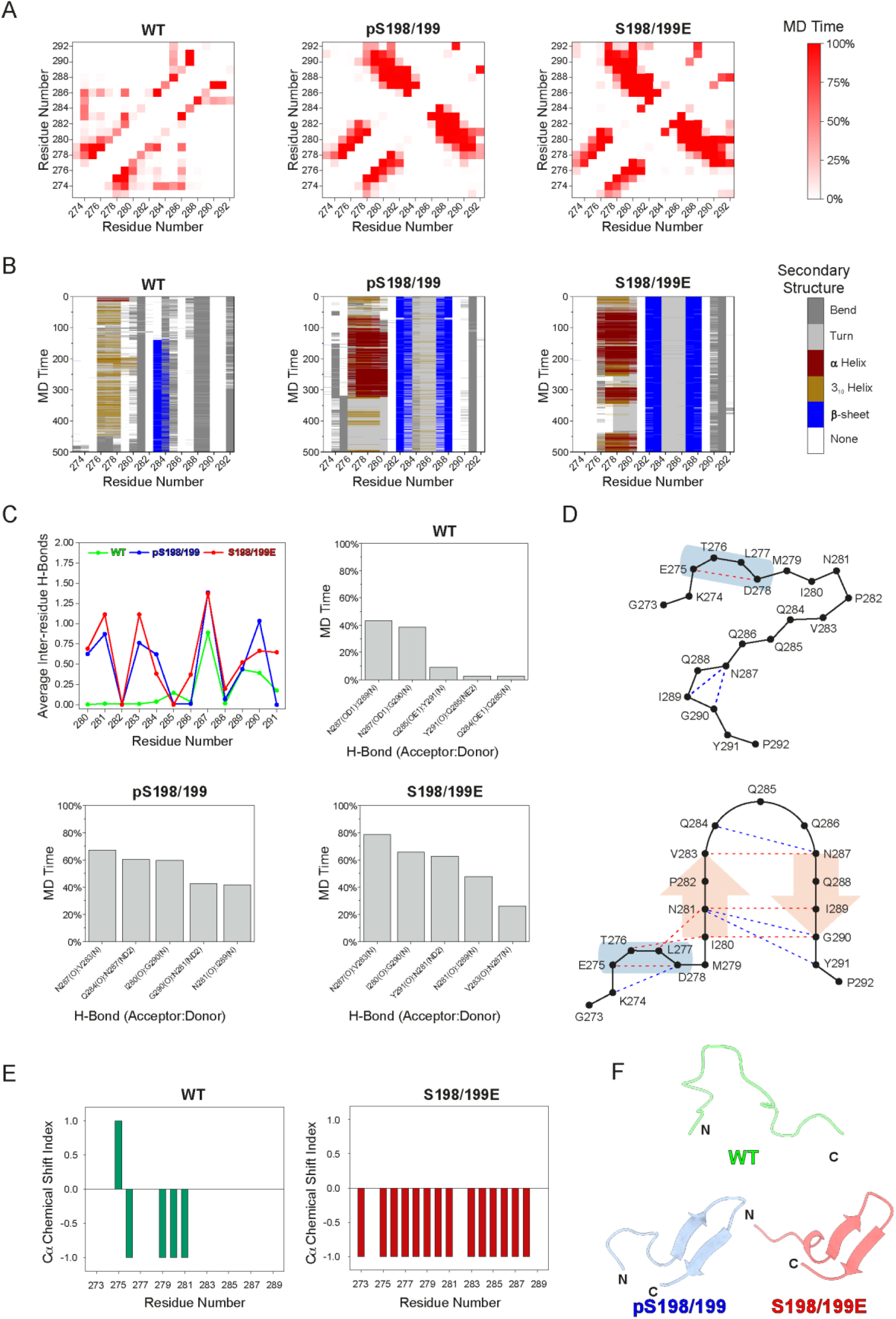
TIA-1 S198/199 phosphorylation and glutamate mutation promote the formation of a β-hairpin motif at the beginning of the PRD. **A)** Inter-residue contact maps obtained from the MD simulations of TIA-1 and covering the first 20 residues of the PRD, where a pattern compatible with a β-hairpin motif is shown for the phosphorylated and phosphomimetic models (colored squares drawing two consecutive diagonals from top to right). A red gradient has been used to indicate the percentage of MD time in which a given contact occurs (cut-off of 6 Å). Only contacts between residues separated by at least 3 residues in sequence have been computed. **B)** Time evolution of the secondary structure of TIA-1 during MD simulations, also centered on the first 20 residues of the PRD. **C)** Comparison of the average number of hydrogen bonds between residues of the predicted β-hairpin motif (280-291) of TIA-1 along MD trajectories (top left panel). Additional bar graphs reveal the top 5 most frequent hydrogen bonds in each model. **D)** Schematic representation of the conformation acquired by the first 20 residues of the PRD in the MD simulations of TIA-1 WT (upper), pS198/199 and S198/199E (lower). The most frequent hydrogen bonds (persistence greater than 25% of the MD time, regardless of occupancy constraints) are shown as dashed lines, colored in red when hydrogen donor and acceptor belong to the protein backbone and in blue when side chain atoms are involved. Secondary structure motifs are also indicated with blue cylinders for helices and red arrows for β-strands. **E)** Chemical Shift Index (CSI) plots derived from the ^13^C_α_ chemical shifts of TIA-1 WT (left panel; BMRB code 18829) and S198/199E (right panel). CSIs were calculated according to ref. 50. In this panel, a minimum of four consecutive “+1” indicates the maximum propensity to form a full α-helix, a minimum of three consecutive “-1” indicates a fully formed β-strand, and “0” indicates disorder. **F)** Representative ribbon structures of the most populated clusters from the MD simulations of TIA-1, showing the β-hairpin region. Clustering of trajectory frames was based on RMSD (residues 191-297).

To confirm the presence of the predicted β-hairpin, NMR experiments (i.e., ^15^N-HSQC, HNCA, and HN(CA)CO) were also acquired for constructs of TIA-1 WT and S198/199E. The ^13^C_α_ chemical shifts of residues G273-I289 (**Supplementary Figure S7**), located in the N-terminal quarter of the PRD, were used to estimate the secondary structure content of both TIA-1 versions^51^. The derived ^13^C_α_ Chemical Shift Indexes suggested that the glutamate mutations induce the folding of the analyzed stretch into a β-strand conformation (**Figure 5E**), supporting that observed in the MD simulations.

### The stability of the identified β-hairpin motif could regulate TIA-1 ability to undergo phase separation

Cross-β interactions have been implicated in the phase transition of a few RBPs into denser fluid states^52–54^, hence the formation of a β-hairpin in TIA-1 PRD could explain the elevated condensation propensity experimentally observed for the phosphomimetic full-length construct. To test this hypothesis, we designed a pair of TIA-1 β-hairpin inhibitory mutants by single substitution of key residues involved in the assembly and stability of the target motif. Tyrosine-to-isoleucine mutation at TIA-1 position 291 (Y291I) would prevent several stabilizing interactions^55–57^ and valine-to-proline mutation at TIA-1 position 283 (V283P) would impair the fundamental V283-N287 hydrogen bond for β-hairpin folding^48–50^. Intriguingly, a valine-to-methionine mutation at position 283 (in TIA-1 isoform a, the equivalent residue is V294) was previously detected in an ALS patient^15^, but the effect of the mutation on the protein remains unknown. Therefore, the full-length TIA-1 β-hairpin mutants Y291I, V283P and V283M were used for computational, *in vitro* and cell-based assays.

First, MD simulations of the aforementioned TIA-1 species were run and analyzed (**Figure 6**). Intramolecular contacts and secondary structure data retrieved from MD trajectories showed that residues of the predicted β-hairpin region tended to be close to each other, especially in V283P and V283M models, but only two residues of the disease-related mutant were consistently assigned in β-strand conformation, namely M283 and N287 (**Figures 6A-B**). However, further analyses did not show a hydrogen bonding pattern consistent with a β-hairpin motif in any TIA-1 model (**Figure 6C**), indicating that none of the mutations lead to β-hairpin formation on their own. Interestingly, the pathological mutation V283M appeared to facilitate the formation of the crucial hydrogen bond ‘closing’ the β-hairpin turn (**Figure 6C**), with occupancy rates slightly higher than in the phosphorylated and phosphomimetic models (**Figure 5C**). However, despite increasing local compactness, this mutation induced a structural configuration substantially different from the β-hairpin found in the simulations of TIA-1 pS198/199 and S198/199E, due to the lack of the necessary hydrogen bonds (**Figures 5C-D and 6C-D**). Thus, according to the computational results, TIA-1 single mutants Y291I and V283P may prevent the formation of a stable β-hairpin in the N-terminal quarter of the PRD, whereas V283M mutation might increase TIA-1 bias towards a specific conformation diverging from that formed under phosphorylation conditions (**Figure 6E**).

**Figure 6.**
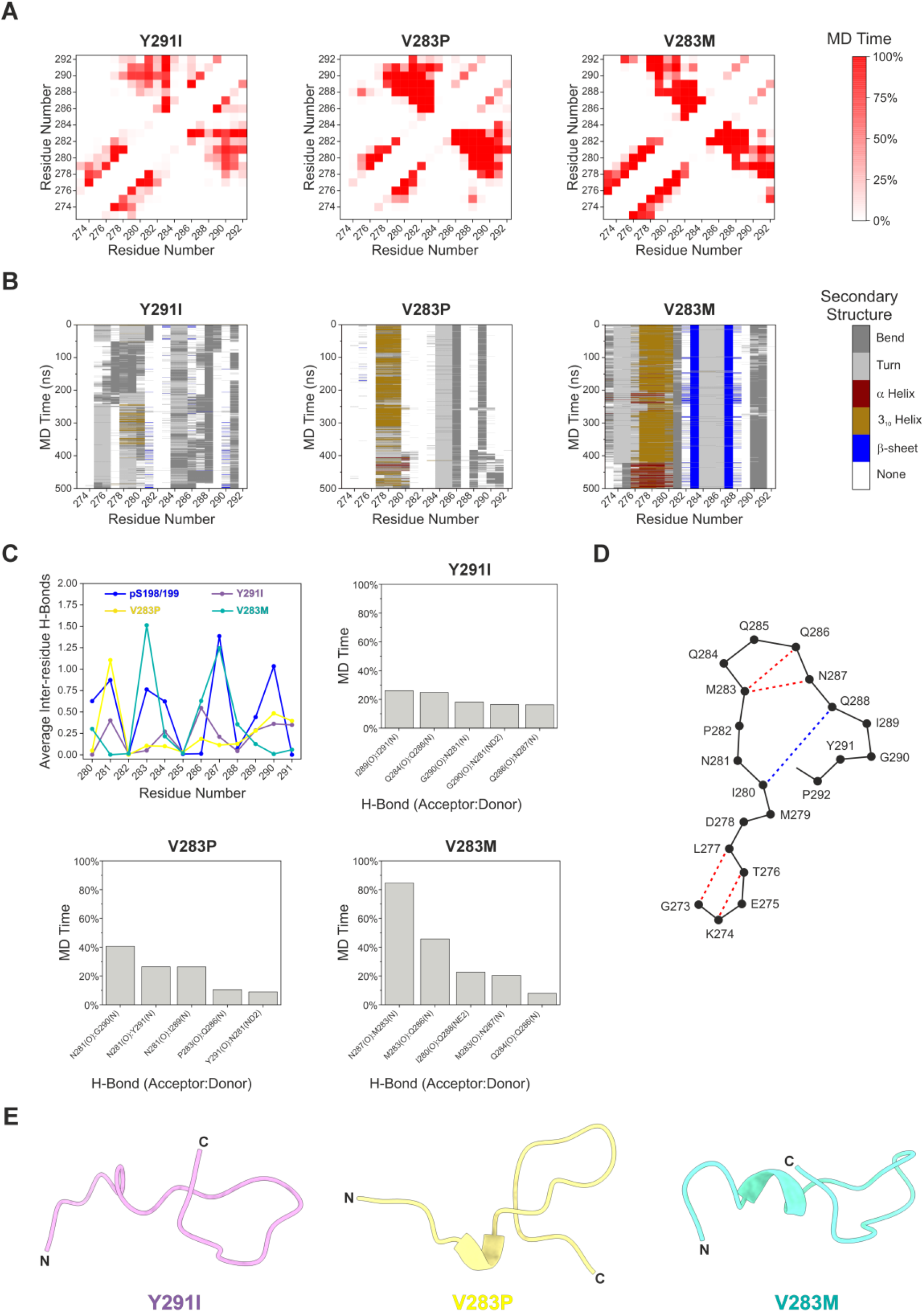
TIA-1 mutations Y291I and V283P hinder β-hairpin formation at the beginning of the PRD, whereas the disease-associated V283M mutation confers this region an aberrant structure. **A)** Inter-residue contact maps obtained from the MD simulations of TIA-1 and covering the first 20 residues of the PRD. A red gradient has been used to indicate the percentage of MD time in which a given contact occurs (cut-off of 6 Å). Only contacts between residues separated by at least 3 residues in sequence have been computed. **B)** Time evolution of the secondary structure of TIA-1 during MD simulations, also centered on the first 20 residues of the PRD. **C)** Comparison of the average number of hydrogen bonds between residues of the predicted β-hairpin motif (280-291) of TIA-1 along MD trajectories (top left panel). Additional bar graphs reveal the top 5 most frequent hydrogen bonds in each model. **D)** Schematic representation of the conformation acquired by the first 20 residues of the PRD in the MD simulation of TIA-1 V283M. The most frequent hydrogen bonds (persistence greater than 25% of the MD time, regardless of occupancy constraints) are shown as dashed lines, colored in red when hydrogen donor and acceptor belong to the protein backbone and in blue when side chain atoms are involved. **E)** Representative ribbon structures of the most populated clusters from the MD simulations of TIA-1, showing the β-hairpin region. Clustering of trajectory frames was based on RMSD (residues 191-297).

Subsequent MD simulations were performed to examine the behavior of TIA-1 bearing one of the β-hairpin mutations along with phosphorylated S198/199. Briefly, analyses revealed that the previously observed phosphorylation-induced β-hairpin assembly is disrupted by the three single mutations tested (**Supplementary Figure S8**). The presence of hydrogen bonds promoting a turn in the ALS-associated TIA-1 mutant (**Figures 6C-D**) was largely decreased in its phosphorylated version (**Supplementary Figure S8C**) and, more importantly, no hydrogen bonding pattern compatible with a β-hairpin motif was detected. In fact, all TIA-1 phosphorylated models presented a more open structure in the putative β-hairpin region as compared with their unphosphorylated counterparts (**Supplementary Figure S8D**). Altogether, our MD data supported the efficacy of TIA-1 β-hairpin inhibitory mutations, even under S198/199 phosphorylation, and predicted that the V283M mutation would also interfere with the formation of a β-hairpin, potentially undermining the pro-LLPS effect of the PTMs under study.

To experimentally corroborate the impact of the aforementioned β-hairpin mutations on SGs assembly, HeLa cells expressing EGFP-tagged TIA-1 full-length Y291I, V283P and V283M constructs were examined by fluorescence microscopy to evaluate the SG formation capacity of the mutant proteins. EGFP-TIA-1 WT and S198/199E were used as controls. Measurements of total cell fluorescence confirmed the presence of similar TIA-1 concentrations in all samples assayed under the same experimental conditions (**Supplementary Figure S1B**). Quantification of the relative area of SGs in unstressed cells did not show meaningful differences between β-hairpin mutants and WT control (**Figure 7A** and **Supplementary Figure S9A**). After arsenite treatment, however, both designed TIA-1 β-hairpin inhibitory mutants (Y291I and V283P) displayed a significantly smaller cellular fraction occupied by SGs compared to WT and the phosphomimetic variant, whereas the pathological TIA-1 V283M appeared midway between the other mutants and control constructs (**Figure 7A** and **Supplementary Figure S9A**). Of note, the percentage of SG-positive cells under stress conditions was considerably lower for TIA-1 Y291I (53%), V283P (73%) and V283M (73%) with respect to WT (85%). To conclude, our experimental results indicated that TIA-1 Y291I, V283P and, to a lesser extent, V283M have a decreased tendency to form SGs.

**Figure 7.**
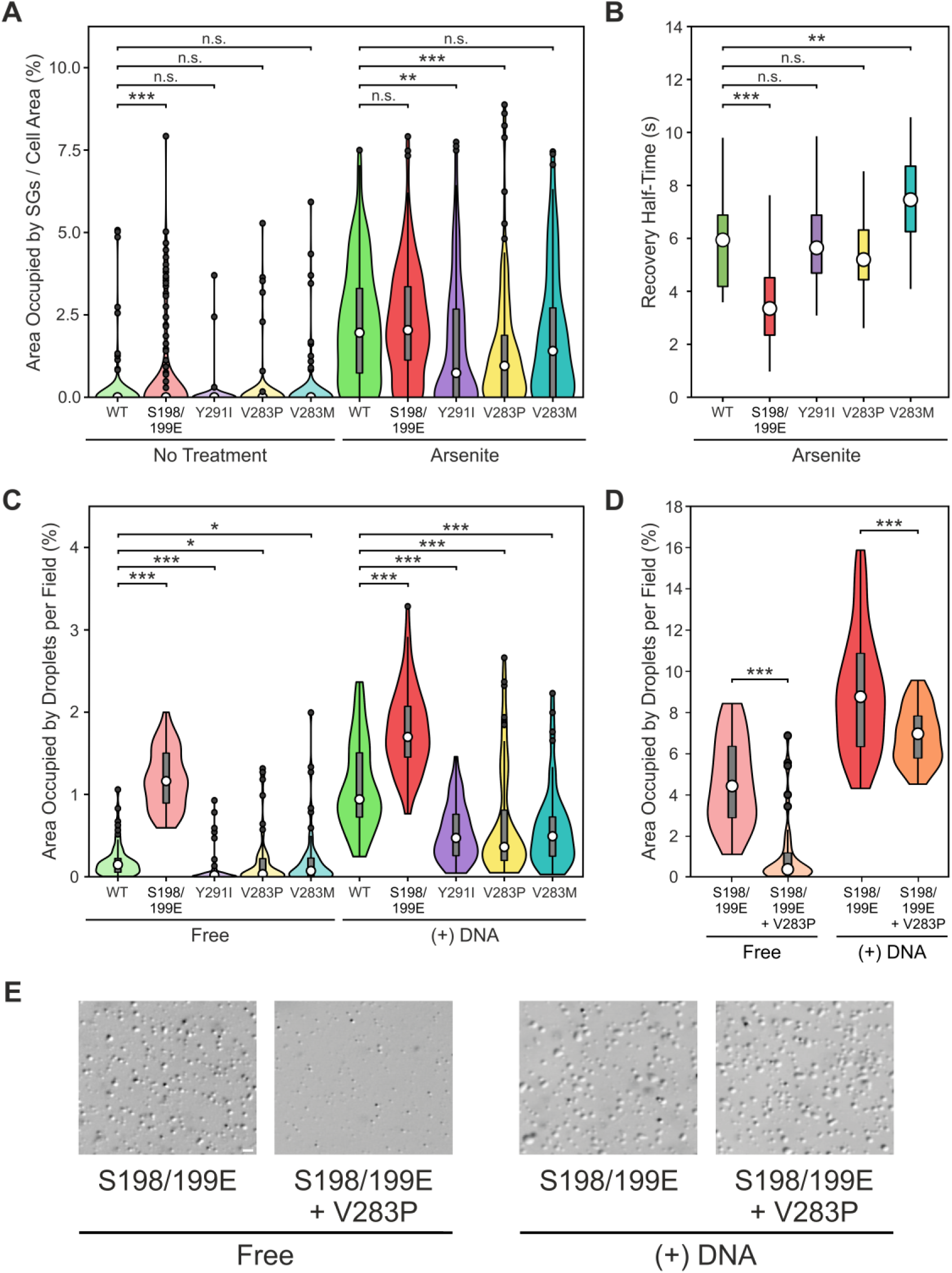
Mutations in the predicted β-hairpin region impinges on TIA-1 condensation properties. **A)** The area of the cell occupied by SGs for each EGFP-TIA-1 construct assayed, expressed by non-treated or arsenite-treated cells, is shown as a violin plot, with the white dot indicating the median; the black dots, the outliers; the box, the interquartile range; and the whiskers extending to the lowest and highest value within 1.5 times, the interquartile range from the hinges. In addition, rotated kernel density plots are given for each dataset to show their distribution. N = 100 cells for each condition. The significance of differences was tested using the Student’s t test. n.s., not significant; \*\**p* < 0.01; \*\*\**p* < 0.001. **B)** FRAP analysis of arsenite-induced SGs containing different EGFP-TIA-1 constructs, showing the half time of fluorescence recovery. White dots indicate the median; boxes for the interquartile range and whiskers extending to the lowest and highest value within 1.5 times for the interquartile range from the hinges. The significance of differences was tested using the Mann Whitney U test. n.s., not significant; \*\**p* < 0.01; \*\*\**p* < 0.001. **C-D)** Droplet formation assays with TIA-1 full-length constructs in the presence and absence of the TC5 ssDNA oligo, showing the area occupied by droplets per field in a violin plot as in (**A**). (**C**) N = 20 fields from 3 replicates. (**D**) N = 13 fields from 3 replicates. The significance of differences was tested using the Mann Whitney U test. n.s., not significant; \**p* < 0.05; \*\*\**p* < 0.001. **E)** Representative bright-field microscopy images of the samples tested in (**D**). Scale bar is 5 μm.

The TIA-1 mutations linked to ALS/FTD studied so far led to low-dynamic and persistent SGs. Specifically, proline-to-hydrophobic residue mutations allowed the disordered PRD structure of TIA-1 to be transformed in β-sheet amyloid-like fibrils, leading to delayed SGs disassembly^15,58^. For that reason, we performed FRAP assays to evaluate the dynamic behavior of TIA-1 WT (control), S198/199E and β-hairpin mutants when assembled into SGs. Arsenite-induced SGs were bleached and then the fluorescence recovery of EGFP-tagged TIA-1 full-length constructs was followed for 44 s. Comparison of recovery half times showed higher turnover rates (i.e., replacement of bleached proteins by fluorescently active proteins) within SGs for phosphomimetic TIA-1, as compared to the other species assayed, including WT (**Figure 7B**). We later confirmed that phosphomimetic TIA-1 possess a reduced RNA-binding affinity (**Supplementary Figure S10**), probably due to the additional negative charge density in RRM3. Of note, the mutations Y291I and V283P did not influence TIA-1 diffusion rate, revealing that the lack of β-hairpin structure would not directly affect the mobility. TIA-1 V283M, on the other hand, exhibited a moderate delay in the recovery half time with respect to WT, which fits well with the pathogenic character of the mutation. Taken together, FRAP assays suggested that S198/199E mutations improve TIA-1 dynamics inside SGs, whereas the ALS-related V283M mutation reduces it.

The LLPS properties of TIA-1 β-hairpin mutants were examined by droplet formation assays in the absence and presence of DNA. The three β-hairpin mutations led to a decrease in the LLPS capacity of the free protein, although more pronounced in the Y291I species (**Figure 7C** and **Supplementary Figure S9B**). This is consistent with the Y291I mutation hindering TIA-1 adoption of appropriate structural configurations for multimerization and subsequent condensation, including but probably not limited to β-hairpin assembly in the initial stretch of the PRD. Furthermore, addition of TC5 ssDNA revealed a reduced LLPS capacity for the three TIA-1 β-hairpin mutants (**Figure 7C** and **Supplementary Figure S9B**). These results are in line with the observed lower propensity to assemble into SGs of these TIA-1 variants (**Figure 7A**), likely due to an impaired ability to establish multivalent homotypic interactions via PRD. Further assays were carried out using the triple mutant construct TIA-1 S198/199E + V283P, aimed at verifying whether the β-hairpin inhibitory mutation can abrogate the pro-LLPS effect of the phosphomimetic mutations. In fact, the V283P substitution caused a strong decline in the droplet formation capacity of phosphomimetic TIA-1 (**Figure 7D**), clearly visible by DIC microscopy (**Figure 7E**), both in the absence and presence of DNA. These results supported the notion that the identified β-hairpin could be responsible for the LLPS-prone behavior observed in TIA-1 S198/199E.

In conclusion, the designed mutations Y291I and V283P notably impaired TIA-1 LLPS and SGs assembly, presumably by precluding the formation and/or destabilizing the β-hairpin motif. On the other hand, the ALS-related mutation V283M may increase SGs persistence despite decreasing TIA-1 condensation propensity, possibly by imposing reduced SGs dynamics that might derive from an aberrant PRD configuration.

## Discussion

TIA-1 double phosphorylation at S198/199 was considered a promising candidate for LLPS modulation given its ‘strategic’ position in RRM3, with both target residues on or around the RNA-interaction surface^29,30^, and spatially close to the initial stretch of the PRD. This suggests the possibility of intramolecular contacts similar to those reported for the FUS family proteins when their PRD serines are phosphorylated^25^. The structure and SG-related function of TIA-1 pS198/199, identified by Tao *et al.*^37^, have been characterized in the present work both computationally and experimentally, unveiling a potential allosteric communication pathway between TIA-1 RRM3 and PRD (**Figure 8**). In essence, we propose that S198/199 phosphorylation in RRM3 allows the formation of a specific network of electrostatic inter-domain interactions that facilitates a disorder-to-order transition in the first quarter of the PRD, creating a β-hairpin motif that promotes TIA-1 LLPS. In contrast, non-phosphorylated TIA-1 would transit between much more conformational states, most of them road-blocks to β-hairpin assembly (**Figure 8** and **Supplementary Figure S11**). As a matter of fact, the phase separation potential of TIA-1 could be attenuated by means of the single mutations Y291I and V283P, designed to prevent the formation and/or destabilize the β-hairpin, even neutralizing the pro-LLPS effect of concurrent phosphomimetic mutations (**Figures 6-7** and **Supplementary Figures S8-S9**). In addition to the increased LLPS propensity, S198/199 phosphorylation endows TIA-1 with greater mobility inside SGs (**Figure 7B**), probably as a consequence of its weakened binding to nucleic acids (**Supplementary Figure S10**). Therefore, considering the pivotal role of TIA-1 as a major constituent and nucleator of SGs, it could be envisaged that S198/199 phosphorylation could lead to more dynamic SGs, easily dismantled upon stress cessation, avoiding their abnormal maturation and pathogenic persistence. Overall, the results presented here support the notion of the β-hairpin-mediated condensation of TIA-1 Ps198/199.

**Figure 8.**
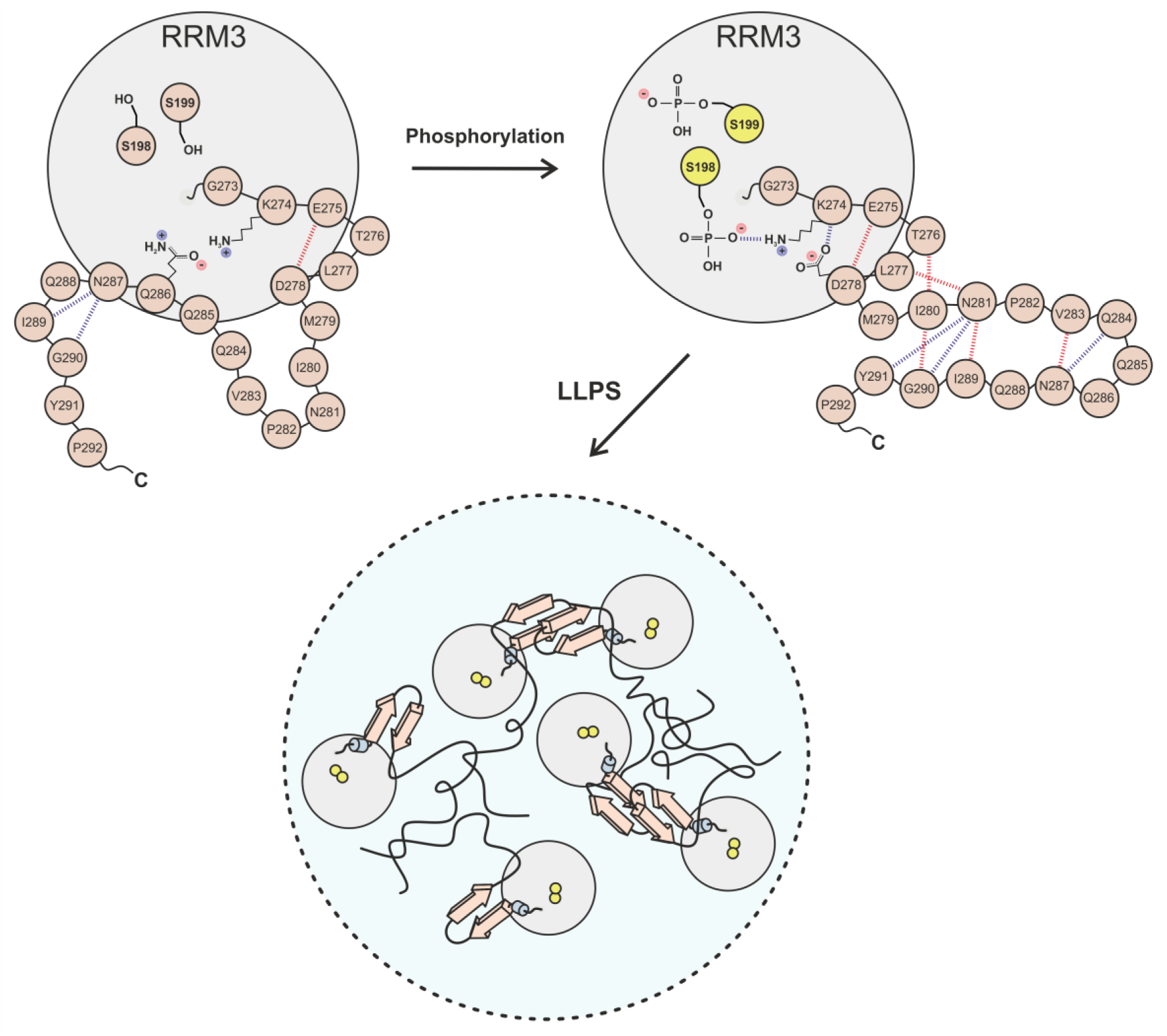
Model of the effect of TIA-1 S198/199 phosphorylation on the disorder-to-order conformational transition leading to protein condensation. The short length and physicochemical nature of unphosphorylated serines restrict the ability of TIA-1 S198/199 to interact with residues at the beginning of the PRD, including K274. The latter residue can associate with nearby glutamines and asparagines of the disordered domain, such as Q286, interactions that suppose road-blocks for the formation of a β-hairpin motif at the beginning of the PRD. In contrast, pS198 can bind strongly to K274 via side chains and limit its freedom to associate with other PRD residues. Indeed, K274 side chain could remain ’trapped’ between pS198 and D278 due to their spatial proximity and charge complementarity. This remodeled network of electrostatic connections would ‘unblock’ the PRD conformational road leading to the adoption of a β-hairpin structure in the N-terminal quarter of the unstructured domain. Then, the ability of TIA-1 to self-associate would be enhanced by stacking of β-hairpin motifs, favoring high local protein concentrations and thus facilitating homotypic intermolecular interactions between other regions of the PRD. In addition, the local rearrangement of the initial stretch of the PRD alone could be enough to alter the global configuration of the domain in a way that promotes multivalence and stability of contacts between sticker regions of different TIA-1 molecules, which would otherwise be entangled in intramolecular interactions. The result of all these changes is a higher propensity of TIA-1 for phase separation into SGs. Note: the most frequent hydrogen bonds (persistence greater than 25% of the MD time, regardless of occupancy constraints) are shown as dashed lines (top panels), colored in red when hydrogen donor and acceptor belong to the protein backbone and in blue when side chain atoms are involved.

The emergence of a β-hairpin motif at the N-terminus of the PRD could influence TIA-1 LLPS by introducing conformational constraints. Basically, the phase separation of proteins is a matter of balance between self-association and solvation, i.e., assembly of protein-rich droplets requires a high degree of cohesion that is achieved by increasing intra- and intermolecular interactions at the expense of solvent exposure^59,60^. In addition, the occurrence of β-turns has been correlated with LLPS for lowering protein desolvation cost^59^. In this sense, the acquisition of a β-hairpin motif involves local compaction of the PRD, which could improve TIA-1 packing efficiency during multimerization and thus favor condensate formation. Another possible consequence of the assembly of the identified β-hairpin is the change in the PRD interaction mode, as a result of a conformational rearrangement that exposes and brings together certain regions of this domain, while conceals and separates others, altering both intra- and intermolecular contacts. According to the stickers-and-spacers model^61^, the sequence of the PRD is composed of a certain number of stretches of variable length (including individual residues) actively implicated in homotypic intermolecular interactions (’stickers’), interspersed with other stretches or residues that influence the PRD binding mode, as well as the structure and dynamics of the droplets (’spacers’).

Therefore, it is conceivable that β-hairpin formation may somehow allow PRD spacers to reach their optimal configuration for maximum cooperativity in the intermolecular interactions of the PRD stickers, thus facilitating TIA-1 LLPS. Interestingly, a parallel mechanism has been proposed for TAR DNA-binding protein of 43 kDa (TDP-43), in which intra-condensate demixing is driven by increased hydrophobic interactions between transient α-helical motifs in its C-terminal domain^62^. These interactions prevent a liquid-to-solid phase transition by suppressing β-sheet aggregation. This analogy suggests that secondary structure-driven rearrangements at the disordered C-terminal domains—whether via β-hairpin formation in TIA-1 or α-helical stabilization in TDP-43— may serve as conserved strategies for modulating the dynamics of RBP condensates. The contribution of Hyman and co-workers further expands this concept by demonstrating that partial local unfolding of the TDP-43 RRM1 domain—specifically exposing cysteine residues—also promotes demixing^62^. This finding aligns well with the micellar model proposed for oligomeric TIA-1 based on solid-state NMR, where RRM1 appears as a dynamically disordered region^63^. Taken together, these insights underscore how conformational flexibility and selective exposure of interaction-prone motifs—whether within structured or disordered regions—can significantly impact phase behavior, suggesting a broader mechanistic framework in which structural plasticity governs condensate regulation across diverse RBPs.

The increase in β-structure content is characteristic of prion and amyloid-β aggregation, where pairs of β-sheet layers are packed together by interdigitating side chains of usually hydrophobic nature, constituting the basic units of fibrils^11,64^. Noteworthy, cross-β interactions are not always conducive to irreversible protein aggregation. For example, the RBPs FUS, hnRNP A1 and hnRNP A2 possess specific stretches or regions of variable length in their IDRs that can adopt different labile cross-β structures and form reversible fibrils^52–54^. These unstable or soft amyloid cores, which would assist LLPS, feature a hydrophilic interface between pairing β-sheet layers enriched in polar residues and coupled together by weaker forces than in classical amyloids. TIA-1 can form both reversible and irreversible amyloid fibrils^15,33,34^. TIA-1 labile fibrils would presumably facilitate the immediate formation of SGs in response to harmful stimuli and their timely clearance upon recovery of cellular homeostasis^65^. Most residues of the phosphorylation-induced TIA-1 β-hairpin have a polar nature, highlighting a glutamine/asparagine (Q/N) rich stretch (284-288), typical of prion and prion-like proteins. Although Q/N-rich regions provide a great tendency to fibrillar aggregation, for many yeast proteins this process is reversible^65,66^. Moreover, the predicted β-hairpin is roughly half hydrophobic and half hydrophilic, and this amphipathic nature most likely limits the strength of potential cross-β interactions. Therefore, this motif might impart instability to TIA-1 amyloid-like fibrils, which would accelerate the formation of SGs while minimizing the risk of conversion into solid inclusions.

Intriguingly, it has recently been reported that the human TIA-1a P362L mutation may induce β-sheet interactions at the C-terminus of the PRD^67,68^, and even enable cross-β interactions by promoting the assembly of a β-hairpin motif specifically at the C-terminal region of the mouse TIA-1 PRD^67^. This ALS-associated mutation grants TIA-1 with an enhanced LLPS and amyloid fibril formation ability *in vitro* and an abnormally high tendency for condensation into low-dynamic and persistent SGs *in cell*^15^. Thus, TIA-1 phase transition into less fluid states seems to be highly sensitive to relatively small changes in the β-sheet content of the PRD, maybe reflecting the delicate equilibrium between TIA-1 functional and pathological fibrillization^65^. The ALS-related mutation V283M appears to be another example of imbalanced fibril formation. This single substitution moderately impaired TIA-1 condensation under stress conditions but also reduced considerably the turnover of SG-resident TIA-1 (**Figure 7** and **Supplementary Figure S9**), suggesting that its pathogenicity could lie in the production of hardened and long-lasting SGs. Strikingly, *in silico* data indicated that the mutation V283M impairs the formation of the β-hairpin despite causing local compaction in unphosphorylated TIA-1 (**Figure 6**), which might influence its conformational dynamics to some degree.

In conclusion, TIA-1 S198/199 phosphorylation emerges as a potential switch mechanism for LLPS regulation, promoting a strong multimerization of this RBP that facilitates the formation of dynamic SGs.

## Methods

### Plasmids and oligonucleotides

Full-length human TIA-1b cloned into the pFRT_TO_eGFP_dest mammalian vector was a gift from Thomas Tuschl (Addgene plasmid #106094). The TIA-1 coding sequence of the above plasmid was cloned into a pETM11 bacterial vector, downstream of a 6xHis-tag, using the oligonucleotides TIA-1 FL-For and TIA-1 FL-Rev. TIA-1 RRM3 25PRD (residues 190-296) sequence, amplified with oligonucleotides TIA-1-RRM3-for2 and TIA-1-25PRD-rev, was cloned into pETM11, replacing the full-length sequence. TIA-1 RRM2,3 (residues 93-274) and TIA-1 RRM2,3 50PRD (residues 93-324) sequences were cloned into a modified pGEX-4T-2 vector (GE Healthcare) bearing a 6xHis-tag and a thrombin cleavage site. Both constructs were inserted downstream of the cleavage sequence using the oligonucleotides TIA-1 RRM2,3-For, TIA-1 RRM2,3-Rev^34^ and TIA-1 RRM2,3 50PRD-Rev. Several different mutations were introduced into the aforementioned plasmids by mutagenic PCR: W147/160/170F, S198/199G, S198/199E, V283M, V283P and Y291I.

All oligonucleotides used are listed in **Supplementary Table S1**. The primers used for molecular cloning as well as TC5 ssDNA^33^ were purchased from StabVida, Eurofins or Sigma-Aldrich. All plasmids generated were checked by sequencing.

### Cell culture and treatment

HeLa cells were cultured in Dulbecco’s modified Eagle’s medium (DMEM) supplemented with 10% heat-inactivated fetal bovine serum (FBS), 2 mM L-glutamine, 100 U/mL penicillin and 100 µg/mL streptomycin, and maintained at 37 °C in a humidified 5% CO_2_ atmosphere (all reactives from Sigma-Aldrich). 80,000-100,000 cells were grown on coverslips in a 24-well plate and plasmids were transfected using Lipofectamine 2000 or 3000 reagent (ThermoFisher) as described by the manufacturer’s instructions, or 1 mg/mL Polyethylenimine hydrochloride (Sigma-Aldrich, in 4-[2-hydroxyethyl]-1-piperazineethanesulphonic acid [HEPES] buffer at pH 8.0). For fluorescence recovery after photobleaching (FRAP) assays, 15,000-30,000 cells were grown on µ-Slides 8 Well Glass Bottom chambered coverslip (ibidi) for 24 h before transfection. Where indicated, cells were treated with 250 μM sodium arsenite (Sigma-Aldrich) for 45 min, approximately 24 h after transfection and just prior fixation or imaging.

### Fluorescence microscopy and stress granules quantification

Quantification of SGs was performed with samples of HeLa cells expressing different EGFP-TIA-1 constructs under basal and oxidative stress conditions. Cells adhered to coverslips were stained with 1 μg/mL Hoechst for nuclei visualization, washed afterwards with phosphate buffer saline (PBS) and fixed with 4% paraformaldehyde (also in PBS) for 10 min at room temperature. Next, cells were washed again with PBS, then immersed in ethanol for 2 min and allowed to air dry. Finally, coverslips were mounted onto slides using Vectashield mounting medium (Vector Laboratories) and sealed with nail polish. Cells were observed using a Leica DM6000 fluorescence microscope (Leica Microsystems) with a 40x objective. The images obtained were processed with ImageJ 1.53n version^69^. Cell contours were manually traced using images of the green fluorescence channel (visualizing EGFP-TIA-1). Nuclei boundaries, on the other hand, were automatically detected using our macro ‘Nuclei Mask’ (see **Supplementary Information** for more details) on images of the blue fluorescence channel (cells stained with Hoechst), discarding those nuclei from cells unsuitable for quantification (mainly non-transfected, but also detached and/or dying cells, and those with poor or excessive fluorescence intensity; see **Supplementary** Figure 1). Regions-of-interest (ROIs) pairs containing the perimeter of each specific cell and its nucleus were used to detect SGs and measure the area occupied (by means of our macro ’SGs Analysis’, see **Supplementary Information**). Boxplots and kernel plots were prepared using R version 4.0.5 (http://www.r-project.org).

### FRAP and cell imaging

HeLa cells were imaged in a laser-scanning confocal microscope TCS SP5 (Leica Microsystems), equipped with temperature incubation and CO_2_ supply. Images were acquired using a 63x/1.4 oil immersion objective. Cells suitable for FRAP experiments were preselected for the EGFP marker of interest. ROIs completely covering the selected SG were bleached with the 488 nm line at 100% laser power. The bleaching protocol was as follows: three pre-bleaching images; five bleaching pulses at 100% laser power and 59 post-bleaching images taken every 0.754 s. Images were collected using the Leica LAS-AF software, recording emitted fluorescence between 487-551 nm.

Fluorescence intensity was measured using ImageJ 1.53n version^69^. Fluorescence mean intensity of ROIs inside bleached SGs and regions around the cell (as a control) were recorded. The mean intensity in these ROIs before bleaching and postbleaching was measured. Background fluorescence in regions outside the cell were also measured. As control, fluorescence mean intensity of nonbleached SGs was recorded. The normalized intensity was plotted over time in the different FRAP experiments (n = 20 cells, from 3-4 independent experiments). Normalized intensity over time was fitted to a single-exponential function (y0+A*(1-exp(-x*k))), and recovery half times were calculated as ln(0.5)/-k, using the Origin v.7.0 software (OriginLab).

### Protein sample preparation

6xHis-tagged TIA-1 RRM2,3 (residues 93-274), TIA-1 RRM2,3 50PRD (residues 93-324), TIA-1 RRM3 25PRD (residues 190-296) and TIA-1 full-length (residues 1-324) constructs were produced in E. coli BL21 (DE3) cells grown at 37°C to an OD_600_ of 0.6-0.8. M9 minimal medium supplemented either with ^15^NH_4_Cl or with ^15^NH_4_Cl and [^13^C]-glucose was used if proteins were intended for NMR measurements, otherwise LB medium was chosen instead. Expression was induced by the addition of 1 mM isopropyl β-D-1-thiogalactopyranoside (IPTG, iNtRON Biotechnology) and cultures were incubated at 30 °C (for truncated TIA-1) or 25 °C (for TIA-1 full-length) during 16-18 h. For extraction of truncated TIA-1 constructs, cell pellets were resuspended in lysis buffer A (20 mM potassium phosphate, 500 mM KCl, pH 7.4) with 20 μg/mL DNase I (Roche), 1 mM phenylmethanesulfonyl fluoride (PMSF, ThermoFisher) and 100 μg/mL lysozyme (ThermoFisher). In the case of TIA-1 full-length species, cell pellets were resuspended in lysis buffer B (50 mM potassium phosphate, 300 mM NaCl, 50 mM Arginine-HCl, 2 mM MgCl_2_, pH 7.0) supplemented with the same additives as before, similarly to ref. 33. Cells were disrupted by sonication (cycles of 30 s at 40% of amplitude, 60 s of rest, 6 min total time, on ice) and cell debris was removed by centrifugation at 28,000 × g (4°C for 45 min). For purification of truncated TIA-1 proteins, the supernatant was loaded onto a Ni-NTA matrix (ThermoFisher), previously equilibrated with lysis buffer A containing 5 mM imidazole, and incubated for 1 h at 4 °C. Recombinant proteins were eluted using a non-continuous imidazole gradient (30-300 mM). Afterwards, they were dialyzed in two steps, at 4 °C overnight, against 5 L of either sodium phosphate buffer or HEPES buffer, whose exact composition and pH varies depending on the downstream experiment. For purification of TIA-1 full-length proteins, the supernatant was also loaded onto a Ni-NTA matrix (ThermoFisher), previously equilibrated with lysis buffer B, and incubated for 1 h at 4 °C. Subsequently, proteins were eluted using a non-continuous imidazole gradient (50-300 mM) in buffer without arginine-HCl. Then, they were dialyzed at 4 °C in two steps, against 5 L of dialysis buffer 1 (20 mM HEPES, 50 mM NaCl, 500 mM arginine-HCl, 2 mM dithiothreitol [DTT], pH 7.4) for 16 h and dialysis buffer 2 (20 mM HEPES, 50 mM NaCl, 500 mM arginine-HCl, 2 mM DTT, 1 mM MgCl_2_, 0.5 mM ethylenediaminetetraacetic acid [EDTA], 5% glycerol, pH 7.4) for 4 h. Purity was further checked by sodium dodecyl sulfate-polyacrylamide gel electrophoresis (SDS-PAGE). Then, samples were concentrated using Amicon Ultra-15 centrifugal filters (Merck-Millipore). Aliquots were concentrated to 100 μM approximately, filtered and stored at -80°C in the final buffer. Proteins were quantified by spectrophotometry at 280 nm using the extinction coefficients estimated with the ProtParam tool (ExPASy server) or, alternatively, by Bradford assay using bovine serum albumin (BSA, ThermoFisher) as standard.

### Turbidity measurements

Turbidity measurements of TIA-1 full-length constructs were performed at a final concentration of 5 μM in 20 mM HEPES, 50 mM NaCl and 1 mM tris(2-carboxyethyl)phosphine (TCEP), at pH 7.4 (droplet buffer), in the presence and absence of 0.5 μM of the multi-site TC5 ssDNA^31^. Independent samples of 100 µL were prepared at room temperature and measured at 385 nm in a 96-well plate with a Varioskan LUX microwell plate reader (ThermoFisher). Each construct was measured at 25 °C every min for 10 min at the indicated ratios. Graphs represent the average of 5 values of turbidity measurements. Assays included 3 replicates with error bars representing the standard deviation (S.D.).

### Droplet formation assay and quantification

Droplet formation of TIA-1 full-length constructs at 10 μM in droplet buffer, with and without TC5 ssDNA at 0.5 μM, was monitored at room temperature by differential interference contrast (DIC) microscopy using a Leica DM6000 fluorescence microscope (Leica Microsystems) with a 40x objective. 60 fields were randomly captured to analyze droplets from 3 independent experiments of each TIA-1 full-length species. DIC images from TIA-1 protein containing S198/199E + V283P mutations (and parental S198/199E as a reference) were recorded using an Olympus FluoView FV3000 LSCM (Olympus) with a 10x objective and 7x zoom. 36 fields were randomly captured to analyze droplets from 3 independent experiments of each TIA-1 full-length species. The images obtained were processed with ImageJ 1.53n version^69^. The total area of the droplets was automatically calculated on images of the bright-field channel using our macro ‘Droplets Analysis’ (see **Supplementary Information**). Results were represented in boxplots and rotated kernel plots as percentages of the area covered by droplets from every field, along with relevant statistical parameters.

### MD simulations and analysis

Molecular models were based on the NMR solution structures of human TIA-1b RRM1 (PBD ID 5O2V^70^) and TIA-1b RRM2,3 (PBD ID 2MJN^29^). The PRD of TIA-1b was added using the built-in module MODELLER^71^ of UCSF Chimera 1.14^72^. Molecular dynamics (MD) computations were carried out on different TIA-1b full-length models (WT, pS198/199 and several mutant variants) using the OpenMM software^73^ in a CUDA platform with the AMBER ff14SB force field. Simulations run under periodic boundary conditions in orthorhombic boxes. Particle Mesh Ewald (PME) electrostatics were set with the Ewald summation cut-off at 10 Å. For each simulation, the system was neutralized with sodium and chlorine counter-ions according to the total charged of the protein, and solvated with optimal 3-charge, 4-point rigid water model (OPC) water molecules. The whole system was subjected to 2,500 steps of energy minimization at 298 K. Langevin thermostat was used to control temperature with a friction coefficient of 1 ps^-1^ and step size of 0.002 ps. Each system was subjected to a total of 700-ns MD simulation and analyzed with the CPPTRAJ module of AmberTools^74^. Radius of gyration (*radgyr* command) and root-mean-square deviation of atomic positions (RSMD, *rms* command) were calculated to check the convergence of the simulations. Accordingly, the last 500 ns of each MD trajectory were further analyzed to evaluate parameters such as specific distances (*distance* command), conformational diversity (hierarchical agglomerative clustering, *hieragglo* command), secondary structure (*secstruct* command), pairwise average electrostatic interaction energy (*pairwise* command, *emapout* keyword) and ‘classical’ hydrogen bonds (NH^…^O) number and lifetime (*hbond* command, *out* and *avgout* keywords). Additionally, intramolecular inter-residue contacts along MD simulations were also analyzed using the Contact Map Explorer Python package^75^, with a cut-off distance of 6 Å, run in the Spyder IDE 3.3.1. Data from analysis were further processed in Origin 2018b (OriginLab) and graphic displays were built in UCSF ChimeraX 1.2.5^76^.

### Fluorescence spectroscopy

Emission spectra were recorded over the wavelength range of 300-500 nm with an excitation wavelength of 295 nm (in order to minimize the contribution from tyrosine fluorescence), using a Perkin-Elmer LS-5 fluorimeter equipped with a water-thermostat cell holder. The slits were set at 5 nm for both excitation and emission, whereas the sample path length was 1 cm. For measurements under native conditions, TIA-1 RRM2,3 W147/160/170F (± S198/199E) at 20 µM was solved into 10 mM sodium phosphate buffer, 50 mM NaCl and 1 mM TCEP, at pH 7.5. For measurements under denaturing conditions, increasing concentrations of urea (6-8 M) were added to the abovementioned buffer and samples of TIA-1 RRM2,3 W147/160/170F at 20 µM were incubated for 1 h at room temperature before recording the spectra. The background signal from the buffer was subtracted. Data reported were the average of 3 replicates.

### NMR spectroscopy

Nuclear magnetic resonance (NMR) experiments with ^15^N-labelled TIA-1 RRM2,3 and TIA-1 RRM2,3 50PRD, both WT and S198/199E, were recorded and monitored at 25 °C by 2D ^1^H-^15^N Heteronuclear Single Quantum Correlation (HSQC) spectra. Additionally, NMR experiments with ^15^N-,^13^C-labelled TIA-1 RRM3 25PRD (residues 190-296) S198/199E were also recorded and monitored at 25 °C by 2D ^1^H-^15^N HSQC, HNCA and HN(CA)CO spectra. All experiments were recorded in a Bruker Avance-III 700 MHz equipped with a 5 mm QCI cryoprobe. Samples of TIA-1 RRM23 and TIA-1 RRM23 50PRD were dialyzed in 10 mM potassium phosphate buffer, 50 mM NaCl and 10 mM DTT, at pH 6.5. Samples of TIA-1 RRM23 25PRD S198/199E were dialyzed in 20 mM potassium phosphate buffer, 50 mM NaCl and 5 mM DTT, at pH 7. To adjust the lock signal, 5% D_2_O was added. Samples of TIA-1 RRM2,3 WT and S198/199E were prepared at 116 or 400 μM, respectively. Samples of TIA-1 RRM2,3 25PRD S198/199E were prepared at 600 or 720 μM, respectively. Samples of TIA-1 RRM2,3 50PRD were prepared at 42 μM for both WT and S198/199E variants. All measurements were taken in 5-mm Shigemi tubes containing samples with a final volume of 350 μL. Water signal was suppressed using the Excitation Sculpting solvent suppression technique. 2D NMR data were acquired and processed using TopSpin 3.5pL7 software (Bruker). 3D NMR data were processed using the software packages NMRPipe/NMRDraw and Topspin (Bruker). Chemical-shift perturbation analyses were performed using the NMRFAM-SPARKY software distribution (National Magnetic Resonance Facility, Madison). The NMR assignment of TIA-1 RRM2,3 and TIA-1 RRM3 18PRD were already available (Biological Magnetic Resonance Bank [BMRB] entries 19735 and 18829, respectively)^29,39^. Weighted average chemical-shift perturbations (Δδ_Avg_) were calculated as previously reported^77^, using the following equation:

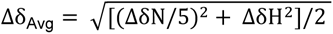

where ΔδN and ΔδH are the chemical shift differences of the amide nitrogen and proton resonances, respectively. Spectra were graphed using CorelDRAW X7.

### CD spectroscopy

Circular dichroism (CD) spectra were recorded in the far-UV range (190-250 nm) on a Jasco J-815 CD spectropolarimeter equipped with a Peltier temperature control system. Samples of TIA-1 RRM2,3 WT and S198/199E at 10 µM in sodium phosphate buffer (pH 7.0) were measured at 20 °C for 20 scans. Samples of TIA-1 RRM2,3 triple tryptophan-to-phenylalanine mutant constructs (±S198/199E) at 10 µM in sodium phosphate buffer (pH 7.5) were measured at 25 °C for 40 scans. All measurements were taken in a 1-mm quartz cuvette. The scanning speed was 100 nm/min. The final spectra were an average of all collected scans.

### FA measurements

Fluorescence anisotropy (FA) assays were performed in 96-well non-binding black plates (Greiner) using a PHERAstar plate reader (BMG Labtech). 5΄-FITC-labelled 10-mer UC1 RNA at 2 nM was incubated at room temperature with either TIA-1 RRM2,3 WT or S198/199E in buffer 10 mM HEPES, 100 mM NaCl and 3% glycerol, at pH 7.0. Data were analyzed using GraphPad Prism and curves were fitted with a single-site specific binding model. Error bars represent error from nonlinear fit.

### Data availability

The authors declare that the main data supporting the findings of this study are contained within the article and its Supplementary Information files or available from the authors upon reasonable request.

## Acknowledgements

This work was supported by Grants PID2024-157414NB-I00 and PID2022-138957NB-I00 funded by MCIN/AEI/10.13039/501100011033 and by “ERDF A way of making Europe”, Ramon Areceś Foundation (LEUCYTO-FRA2025), Andalusian Government (BIO-198) and the National Health and Medical Research Council of Australia [APP1105801] awarded to J.A.W, the Australian Research Council [DP200102737] awarded to J.A.W. and F.E.L. A.V.-C. was awarded a PhD fellowship from the Spanish Ministry of Education, Culture and Sport (FPU016/01513), L. C.G. supported by the Andalusian Government (PAIDI-Doctor 2020 DOC_00796) and A.B.U was recognized with a Technical Support Staff contract (PTA2023-022984-I) and a Juan de la Cierva contract (JDC2023-050813-I), both funded by the Spanish Ministry of Science and Innovation. E.T. was supported by CIBERNED (ISCIII) contract. We thank the staff at the NMR facility at the Research, Technology and Innovation center of the University of Seville (CITIUS), and the BIO-MS laboratory (UPO), the microscopy, chromatography and centrifuges services at the Institute of Plant Biochemistry and Photosynthesis (IBVF), the Biomolecular Interaction Platform and (cicCartuja, Seville), and the microscopy service of CABIMER and the Proteomics Facility at the Institute of Biomedicine of Seville (IBiS).

## Author contributions

A.V.C., L.C.G., A.B.U.M., R.F.C. and I.D.M. designed the study. A.V.C., L.C.G., R.L.G.A. and E.T. generated the strains, purified the proteins and performed the experiments. A.V.C., A.B.U.M., S.M.G.M. and I.D.M. performed the MDs and NMR experiments. S.W. and F.L. performed the fluorescence anisotropy assays. I.D.M, M.A.R., R.F.C. and J.A.W. acquired funding. I.D.M, M.A.R., R.F.C. and J.A.W. supervised the work. A.V.C, L.C.G., A.B.U.M. and I.D.M wrote the manuscript, with input from all other authors.

## References

1. Yamasaki, S. & Anderson, P. Reprogramming mRNA translation during stress. Curr. Opin. Cell Biol. 20, 222–226 (2008).

2. Buchan, J. R., Yoon, J. H. & Parker, R. Stress-specific composition, assembly and kinetics of stress granules in Saccharomyces cerevisiae. J. Cell Sci. 124, 228–239 (2011).

3. Fefilova, A. S., Fonin, A. V., Vishnyakov, I. E., Kuznetsova, I. M. & Turoverov, K. K. Stress-Induced Membraneless Organelles in Eukaryotes and Prokaryotes: Bird’s-Eye View. Int. J. Mol. Sci. 23, 5010 (2022).

4. Aulas, A. & Vande Velde, C. Alterations in stress granule dynamics driven by TDP-43 and FUS: a link to pathological inclusions in ALS? Front. Cell. Neurosci. 9, 423 (2015).

5. Mahboubi, H. & Stochaj, U. Cytoplasmic stress granules: Dynamic modulators of cell signaling and disease. Biochim. Biophys. Acta - Mol. Basis Dis. 1863, 884–895 (2017).

6. Protter, D. S. W. & Parker, R. Principles and Properties of Stress Granules. Trends Cell Biol. 26, 668–679 (2016).

7. Kedersha, N., Ivanov, P. & Anderson, P. Stress granules and cell signaling: more than just a passing phase? Trends Biochem. Sci. 38, 494–506 (2013).

8. Kedersha, N. et al. Dynamic shuttling of TIA-1 accompanies the recruitment of mRNA to mammalian stress granules. J. Cell Biol. 151, 1257–1268 (2000).

9. Panas, M. D., Ivanov, P. & Anderson, P. Mechanistic insights into mammalian stress granule dynamics. J. Cell Biol. 215, 313–323 (2016).

10. Ivanov, P., Kedersha, N. & Anderson, P. Stress Granules and Processing Bodies in Translational Control. Cold Spring Harb. Perspect. Biol. 11, a032813 (2019).

11. Babinchak, W. M. & Surewicz, W. K. Liquid-Liquid Phase Separation and Its Mechanistic Role in Pathological Protein Aggregation. J. Mol. Biol. 432, 1910–1925 (2020).

12. Dudman, J. & Qi, X. Stress Granule Dysregulation in Amyotrophic Lateral Sclerosis. Front. Cell. Neurosci. 14, 598517 (2020).

13. Nedelsky, N. B. & Taylor, J. P. Pathological phase transitions in ALS-FTD impair dynamic RNA-protein granules. RNA 28, 97–113 (2022).

14. Wang, J., Gan, Y., Cao, J., Dong, X. & Ouyang, W. Pathophysiology of stress granules: An emerging link to diseases (Review). Int. J. Mol. Med. 49, 44 (2022).

15. Mackenzie, I. R. et al. TIA1 Mutations in Amyotrophic Lateral Sclerosis and Frontotemporal Dementia Promote Phase Separation and Alter Stress Granule Dynamics. Neuron 95, 808–816.e9 (2017).

16. Ohn, T. & Anderson, P. The role of posttranslational modifications in the assembly of stress granules. Wiley Interdiscip. Rev. RNA 1, 486–493 (2010).

17. García-Mauriño, S. M. et al. RNA Binding Protein Regulation and Cross-Talk in the Control of AU-rich mRNA Fate. Front. Mol. Biosci. 4, 71 (2017).

18. Velázquez-Cruz, A., Baños-Jaime, B., Díaz-Quintana, A., De la Rosa, M. A. & Díaz-Moreno, I. Post-translational Control of RNA-Binding Proteins and Disease-Related Dysregulation. Front. Mol. Biosci. 8, 658852 (2021).

19. Arenas, A. et al. Lysine acetylation regulates the RNA binding, subcellular localization and inclusion formation of FUS. Hum. Mol. Genet. 29, 2684–2697 (2020).

20. Guillén-Boixet, J. et al. RNA-Induced Conformational Switching and Clustering of G3BP Drive Stress Granule Assembly by Condensation. Cell 181, 346–361.e17 (2020).

21. Yang, P. et al. G3BP1 Is a Tunable Switch that Triggers Phase Separation to Assemble Stress Granules. Cell 181, 325–345.e28 (2020).

22. Tsai, W.-C. et al. Arginine Demethylation of G3BP1 Promotes Stress Granule Assembly. J. Biol. Chem. 291, 22671–22685 (2016).

23. Qamar, S. et al. FUS Phase Separation Is Modulated by a Molecular Chaperone and Methylation of Arginine Cation-pi Interactions. Cell 173, 720–734.e15 (2018).

24. Hofweber, M. et al. Phase Separation of FUS Is Suppressed by Its Nuclear Import Receptor and Arginine Methylation. Cell 173, 706–719.e13 (2018).

25. Wang, J. et al. A Molecular Grammar Governing the Driving Forces for Phase Separation of Prion-like RNA Binding Proteins. Cell 174, 688–699.e16 (2018).

26. Andrusiak, M. G. et al. Inhibition of Axon Regeneration by Liquid-like TIAR-2 Granules. Neuron 104, 290–304.e8 (2019).

27. Förch, P. et al. The apoptosis-promoting factor TIA-1 is a regulator of alternative pre-mRNA splicing. Mol. Cell 6, 1089–1098 (2000).

28. Cruz-Gallardo, I. et al. The binding of TIA-1 to RNA C-rich sequences is driven by its C-terminal RRM domain. RNA Biol. 11, 766–776 (2014).

29. Wang, I. et al. Structure, dynamics and RNA binding of the multi-domain splicing factor TIA-1. Nucleic Acids Res. 42, 5949–5966 (2014).

30. Waris, S. et al. TIA-1 RRM23 binding and recognition of target oligonucleotides. Nucleic Acids Res. 45, 4944–4957 (2017).

31. Suswman, E. A., Li, Y. Y., Mahtani, H. & King, P. H. Novel DNA-binding properties of the RNA-binding protein TIAR. Nucleic Acids Res. 33, 4507–4518 (2005).

32. Gilks, N. et al. Stress granule assembly is mediated by prion-like aggregation of TIA-1. Mol. Biol. Cell 15, 5383–5398 (2004).

33. Loughlin, F. E. et al. Tandem RNA binding sites induce self-association of the stress granule marker protein TIA-1. Nucleic Acids Res. 49, 2403–2417 (2021).

34. West, D. L. et al. Regulation of TIA-1 Condensates: Zn(2+) and RGG Motifs Promote Nucleic Acid Driven LLPS and Inhibit Irreversible Aggregation. Front. Mol. Biosci. 9, 960806 (2022).

35. Corrales-Guerrero, L. & Díaz-Moreno, I. Deciphering the role of Zn(2+)-binding histidines from TIA-1 on the assembly and dynamics of stress granules. BioFactors 50, 750–755 (2024).

36. Hornbeck, P. V. et al. PhosphoSitePlus, 2014: mutations, PTMs and recalibrations. Nucleic Acids Res. 43, D512–520 (2015).

37. Tao, W. A. et al. Quantitative phosphoproteome analysis using a dendrimer conjugation chemistry and tandem mass spectrometry. Nat. Methods 2, 591–598 (2005).

38. Aroca, A., Díaz-Quintana, A. & Díaz-Moreno, I. A structural insight into the C-terminal RNA recognition motifs of T-cell intracellular antigen-1 protein. FEBS Lett. 585, 2958–2964 (2011).

39. Cruz-Gallardo, I., Aroca, A., Persson, C., Karlsson, B. G. & Díaz-Moreno, I. RNA binding of T-cell intracellular antigen-1 (TIA-1) C-terminal RNA recognition motif is modified by pH conditions. J. Biol. Chem. 288, 25986–25994 (2013).

40. Vivian, J. T. & Callis, P. R. Mechanisms of tryptophan fluorescence shifts in proteins. Biophys. J. 80, 2093–2109 (2001).

41. Royer, C. A. Probing protein folding and conformational transitions with fluorescence. Chem. Rev. 106, 1769–1784 (2006).

42. Ghisaidoobe, A. B. & Chung, S. J. Intrinsic tryptophan fluorescence in the detection and analysis of proteins: a focus on Forster resonance energy transfer techniques. Int. J. Mol. Sci. 15, 22518–22538.

43. Chen, Y., Liu, B., Yu, H.-T. & Barkley, M. D. The Peptide Bond Quenches Indole Fluorescence. J. Am. Chem. Soc. 118, 9271–9278 (1996).

44. Chen, J., Callis, P. R. & King, J. Mechanism of the very efficient quenching of tryptophan fluorescence in human gamma D- and gamma S-crystallins: the gamma-crystallin fold may have evolved to protect tryptophan residues from ultraviolet photodamage. Biochemistry 48, 3708–3716 (2009).

45. Martin, E. W. et al. Interplay of folded domains and the disordered low-complexity domain in mediating hnRNPA1 phase separation. Nucleic Acids Res. 49, 2931–2945 (2021).

46. Milner-White, E. J. & Poet, R. Four classes of beta-hairpins in proteins. Biochem. J. 240, 289– 292 (1986).

47. Blanco, F., Ramirez-Alvarado, M. & Serrano, L. Formation and stability of beta-hairpin structures in polypeptides. Curr. Opin. Cell Biol. 8, 107–111 (1998).

48. Muñoz, V., Henry, E. R., Hofrichter, J. & Eaton, W. A. A statistical mechanical model for beta-hairpin kinetics. Proc. Natl. Acad. Sci. U. S. A. 95, 5872–5879 (1998).

49. Lindorff-Larsen, K., Piana, S., Dror, R. O. & Shaw, D. E. How fast-folding proteins fold. Science 334, 517–520 (2011).

50. Enemark, S. & Rajagopalan, R. Turn-directed folding dynamics of beta-hairpin-forming de novo decapeptide Chignolin. Phys. Chem. Chem. Phys. 14, 12442–12450 (2012).

51. Wishart, D. S. & Sykes, B. D. The 13C chemical-shift index: a simple method for the identification of protein secondary structure using 13C chemical-shift data. J. Biomol. NMR 4, 171–180 (1994).

52. Luo, F. et al. Atomic structures of FUS LC domain segments reveal bases for reversible amyloid fibril formation. Nat. Struct. Mol. Biol. 25, 341–346 (2018).

53. Gui, X. et al. Structural basis for reversible amyloids of hnRNPA1 elucidates their role in stress granule assembly. Nat. Commun. 10, 2006 (2019).

54. Kato, M. & McKnight, S. L. The low-complexity domain of the FUS RNA binding protein self-assembles via the mutually exclusive use of two distinct cross-beta cores. Proc. Natl. Acad. Sci. U. S. A. 118, e2114412118 (2021).

55. Waters, M. L. Aromatic interactions in peptides: impact on structure and function. Biopolymers 76, 435–445 (2004).

56. Valley, C. C. et al. The methionine-aromatic motif plays a unique role in stabilizing protein structure. J. Biol. Chem. 287, 34979–34991 (2012).

57. Mahalakshmi, R. Aromatic interactions in beta-hairpin scaffold stability: A historical perspective. Arch. Biochem. Biophys. 661, 39–49 (2019).

58. Ding, X., Gu, S., Xue, S. & Luo, S. Z. Disease-associated mutations affect TIA1 phase separation and aggregation in a proline-dependent manner. Brain Res. 1768, 147589 (2021).

59. Paiz, E. A. et al. Beta turn propensity and a model polymer scaling exponent identify intrinsically disordered phase-separating proteins. J. Biol. Chem. 297, 101343 (2021).

60. Ibrahim, A. Y. et al. Intrinsically disordered regions that drive phase separation form a robustly distinct protein class. J. Biol. Chem. 299, 102801 (2023).

61. Ginell, G. M. & Holehouse, A. S. An Introduction to the Stickers-and-Spacers Framework as Applied to Biomolecular Condensates. Methods Mol. Biol. 2563, 95–116 (2023).

62. Yan, X. et al. Intra-condensate demixing of TDP-43 inside stress granules generates pathological aggregates. Cell 188, 4123–4140.e18 (2025).

63. Fritzsching, K. J. et al. Micellar TIA1 with folded RNA binding domains as a model for reversible stress granule formation. Proc. Natl. Acad. Sci. U. S. A. 117, 31832–31837 (2020).

64. Ono, K. & Watanabe-Nakayama, T. Aggregation and structure of amyloid beta-protein. Neurochem. Int. 151, 105208 (2021).

65. Furukawa, Y. & Nukina, N. Functional diversity of protein fibrillar aggregates from physiology to RNA granules to neurodegenerative diseases. Biochim. Biophys. Acta 1832, 1271–1278 (2013).

66. Cereghetti, G., Saad, S., Dechant, R. & Peter, M. Reversible, functional amyloids: towards an understanding of their regulation in yeast and humans. Cell Cycle 17, 1545–1558 (2018).

67. Sekiyama, N. et al. ALS mutations in the TIA-1 prion-like domain trigger highly condensed pathogenic structures. Proc. Natl. Acad. Sci. U. S. A. 119, e2122523119 (2022).

68. Wittmer, Y. et al. Liquid Droplet Aging and Seeded Fibril Formation of the Cytotoxic Granule Associated RNA Binding Protein TIA1 Low Complexity Domain. J. Am. Chem. Soc. 145, 1580– 1592 (2023).

69. Schneider, C. A., Rasband, W. S. & Eliceiri, K. W. NIH Image to ImageJ: 25 years of image analysis. Nat. Methods 9, 671–675 (2012).

70. Sonntag, M., et al. Segmental, Domain-Selective Perdeuteration and Small-Angle Neutron Scattering for Structural Analysis of Multi-Domain Proteins. Angew. Chem. Int. Ed. 56, 9322–9325 (2017).

71. Webb, B. & Sali, A. Comparative Protein Structure Modeling Using MODELLER. Curr. Protoc. Protein Sci. 86, 2.9.1–2.9.37 (2016).

72. Pettersen, E. F. et al. UCSF Chimera--a visualization system for exploratory research and analysis. J. Comput. Chem. 25, 1605–1612 (2004).

73. Eastman, P. et al. OpenMM 7: Rapid development of high performance algorithms for molecular dynamics. PLoS Comput. Biol. 13, e1005659 (2017).

74. Roe, D. R. & Cheatham, T. E. 3rd. PTRAJ and CPPTRAJ: Software for Processing and Analysis of Molecular Dynamics Trajectory Data. J. Chem. Theory Comput. 9, 3084–3095 (2013).

75. 75. Swenson, D. W. H. & Roet, S. Contact Map Explorer (Version 2.1) [Source code] https://github.com/dwhswenson/contact_map. (2017).

76. Pettersen, E. F. et al. UCSF ChimeraX: Structure visualization for researchers, educators, and developers. Protein Sci. 30, 70–82 (2021).

77. Díaz-Moreno, I. et al. Orientation of the central domains of KSRP and its implications for the interaction with the RNA targets. Nucleic Acids Res. 38, 5193–5205 (2010).

