## Supplementary Information for "Allosteric modulation of TIA-1 phase separation by double serine phosphorylation"

Alejandro Velázquez-Cruz[1], Laura Corrales-Guerrero[1,‡], Ana B. Uceda-Mayo[1,#], Emanuela Tumini[2], Sofía M. García-Mauriño[1], Saboora Waris[3,†], Rafael L. Giner-Arroyo[1], Fionna E. Loughlin[3], Rafael Fernández-Chacón[2], Jacqueline A. Wilce[3], Miguel A. De la Rosa[1] and Irene Díaz-Moreno[1,*]

[1] Institute for Chemical Research — Centro de Investigaciones Científicas Isla de la Cartuja (cicCartuja), Universidad de Sevilla – Consejo Superior de Investigaciones Científicas (CSIC), Seville, Spain.
[2] Institute of Biomedicine of Seville (IBiS, University Hospital Virgen del Rocío – CSIC – University of Seville, Department of Medical Physiology and Biophysics, School of Medicine, and CIBERNED ISCIII, Seville, Spain
[3] Monash Biomedicine Discovery Institute and Department of Biochemistry and Molecular Biology, Monash University, Clayton, VIC, Australia.

* To whom correspondence should be addressed.

‡ Present address: Institute of Plant Biochemistry and Photosynthesis, — Centro de Investigaciones Científicas Isla de la Cartuja (cicCartuja), Universidad de Sevilla – Consejo Superior de Investigaciones Científicas (CSIC), Seville, Spain.

### Present address: Department of Chemistry, University of the Balearic Islands – Health Research Institute of the Balearic Islands (IdISBa) – Institut Universitari d'Investigació en Ciències de la Salut (IUNICS), Palma de Mallorca, Spain.

† Present address: Maroof Clinical Trial Unit, Research Department, Maroof International Hospital Islamabad, Pakistan.

**This PDF includes:**

- Supplementary Figures S1 to S11
- Supplementary Table S1
- ImageJ macros

### Supplementary Information

#### Supplementary Tables

##### Table S1. Oligonucleotides used in this work.

| Name | Sequence (5' to 3') | Reference |
|---|---|---|
| TIA-1-W147/160/170F-For | GATGGGTGGCCAGtttCTTGGTGGAAGACAAATCAGAACTAACtttGCAA | Aroca et al., 2011 |
| TIA-1-W147/160/170F-Rev | CAAGaaaCTGGCCACCCATCTGTTGAATGGCGTTTTCAGCATCaaaTTTGT | Aroca et al., 2011 |
| TC1 | TTTTTACTCC | Loughlin et al., 2021 |
| TC5 | TTTTTACTCCAATTTTTACTCCAATTTTTACTCCAATTTTTACTCCAATTTTTACTCC | Loughlin et al., 2021 |
| TIA-1 RRM2,3-For | GCGTGGATCCCCAGGAATTCCCAATCATTTCCATGTCTTTGTTGGTG | West et al., 2022 |
| TIA-1 RRM2,3-Rev | CAGTCACGATGCGGCCGCTTATTTGCCCCAATAGCATTTCACAAC | West et al., 2022 |
| TIA-1 FL-For | GCGGATCCGAATTCATGGAGGACGAGATGCCCAAG | This study |
| TIA-1 FL-Rev | TCGAGTGCGGCCGCTCATCACTGGGTTTCATAC | This study |
| TIA-1 RRM2,3 50PRD-Rev | CAGTCACGATGCGGCCGCTTA-ATACATTCCATATGCAGGAACTTGC | This study |
| TIA-1-S198/199E-For | ATCAGgaggagCCAAGCAACTGTACTGTATACTGTGGAGG | This study |
| TIA-1-S198/199E-Rev | TTGGctcctcCTGATTTACAACCTCATCATATGATAGCTG | This study |
| TIA-1-S198/199G-For | ATCAGggtggtCCAAGCAACTGTACTGTATACTGTGGAGG | This study |
| TIA-1-S198/199G-Rev | TTGGaccaccCTGATTTACAACCTCATCATATGATAGCTG | This study |
| TIA-1-V283M-For | AATCCCatgCAACAGCAGAATCAAATTGG | This study |
| TIA-1-V283M-Rev | TGCTGTTGtacGGGATTTATCATATCAAGAG | This study |
| TIA-1-V283P-For | AATCCCccgCAACAGCAGAATCAAATTGG | This study |
| TIA-1-V283P-Rev | TGCTGTTGGGcggGATTTATCATATCAAGAG | This study |
| TIA-1-Y291I-For | ATTGGAattCCCCAACCTTATGGCCAGTG | This study |
| TIA-1-Y291I-Rev | TTGGGGaatTCCAATTTGATTCTGCTGTTG | This study |
| TIA-1-25PRD-rev | GAGTGCGGCCGCTCAGCCATAAGGTTGGGGATATCCA | This study |
| TIA-1-RRM3-for2 | AGCGGATCCGAATTCTCATATGATGAGGTTGTAAATCAG | This study |

#### Supplementary Figures

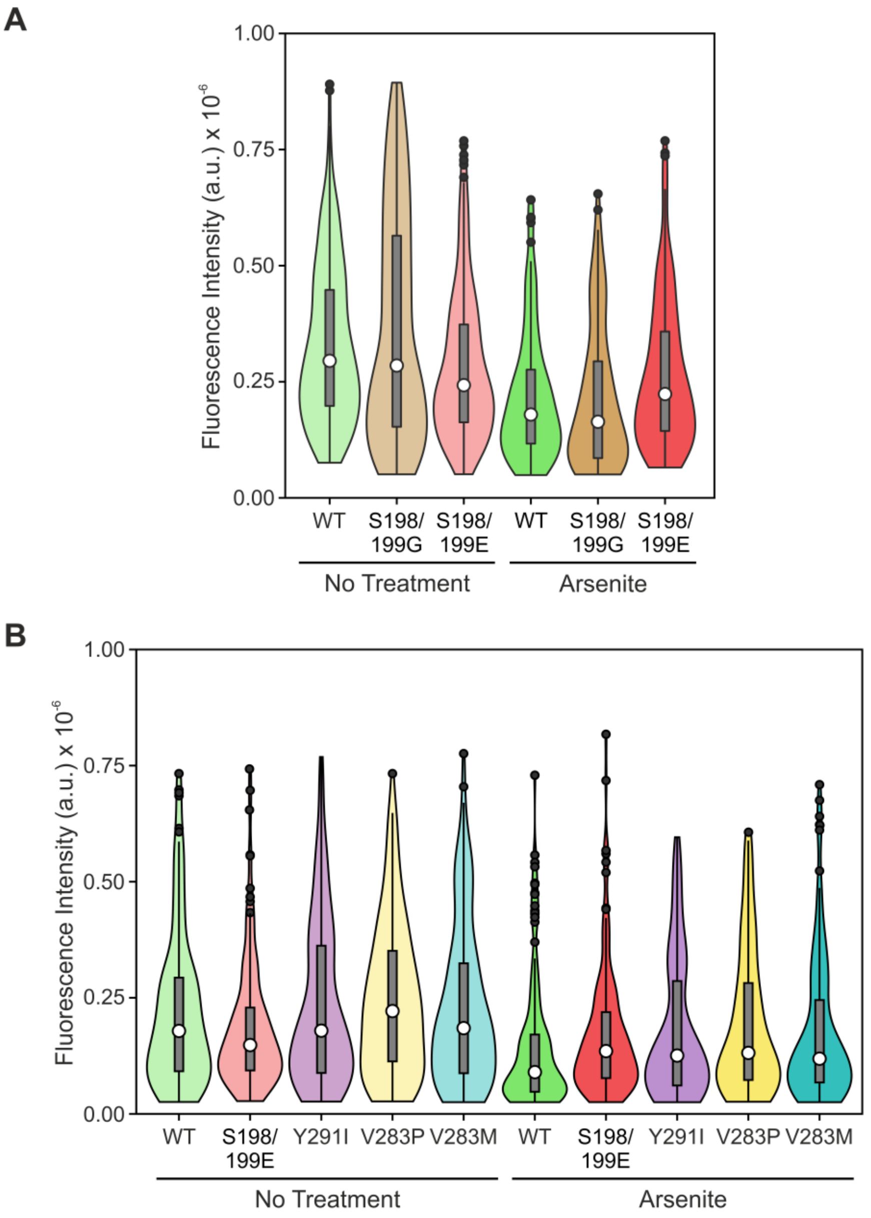


**Supplementary Figure S1. Quantification of EGFP-TIA-1 expression levels in the fluorescence microscopy experiments.** Fluorescence intensity measurements of EGFP-TIA-1 constructs assayed in two independent sets of experiments (**A** and **B**), expressed by non-treated or arsenite-treated cells, are shown as violin plots, with the white dot indicating the median; the black dots, the outliers; the box, the interquartile range; and the whiskers extending to the lowest and highest value within 1.5 times, the interquartile range from the hinges. In addition, rotated kernel density plots are given for each dataset to show their distribution. (**A**) N = 300 cells for basal conditions and 150 cells for stress conditions. (**B**) N = 100 cells for each condition.

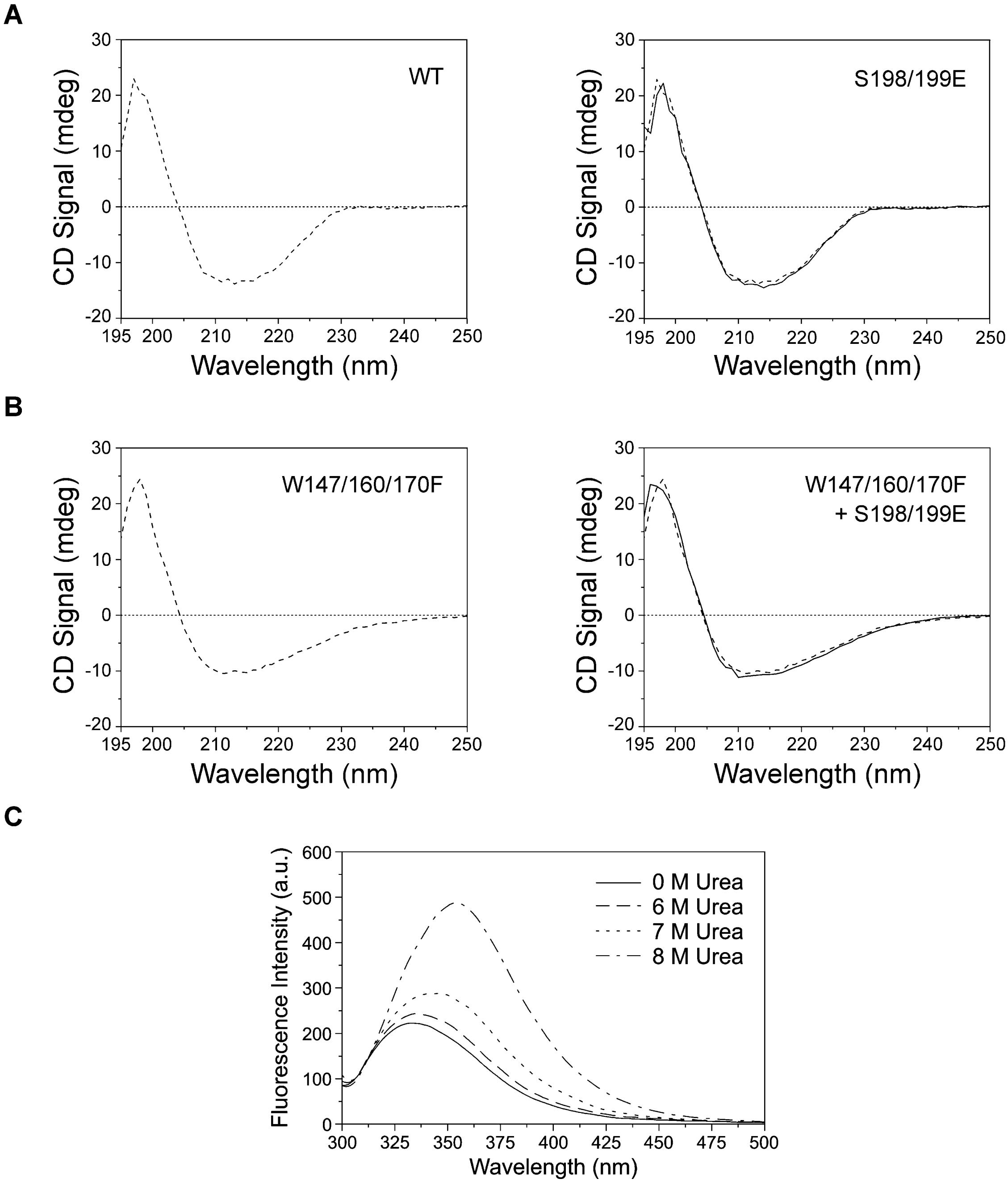


**Supplementary Figure S2. Secondary structure analysis and fluorescence spectroscopy measurements. A-B)** Far-UV circular dichroism spectra of TIA-1 RRM2,3 WT and S198/199E (**A**), and their triple tryptophan-to-phenylalanine mutant constructs (**B**). **C)** Fluorescence titration of TIA-1 RRM2,3 W147/160/170F with increasing concentrations of urea.

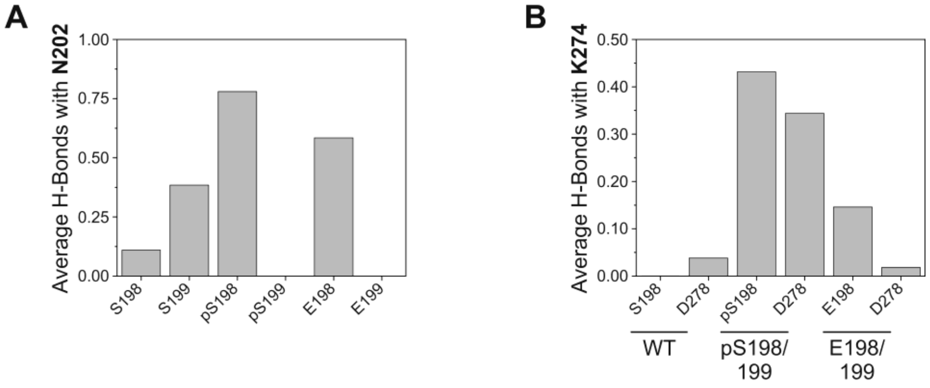


**Supplementary Figure S3. Hydrogen bond analyses of TIA-1 WT, pS198/199 and S198/199E MD simulations. A-B)** Bar graphs comparing the average number of hydrogen bonds between residues 198/199 and N202 (**A**), as well as between K274 and residues 198 and D278 (**B**), during TIA-1 MD trajectories.

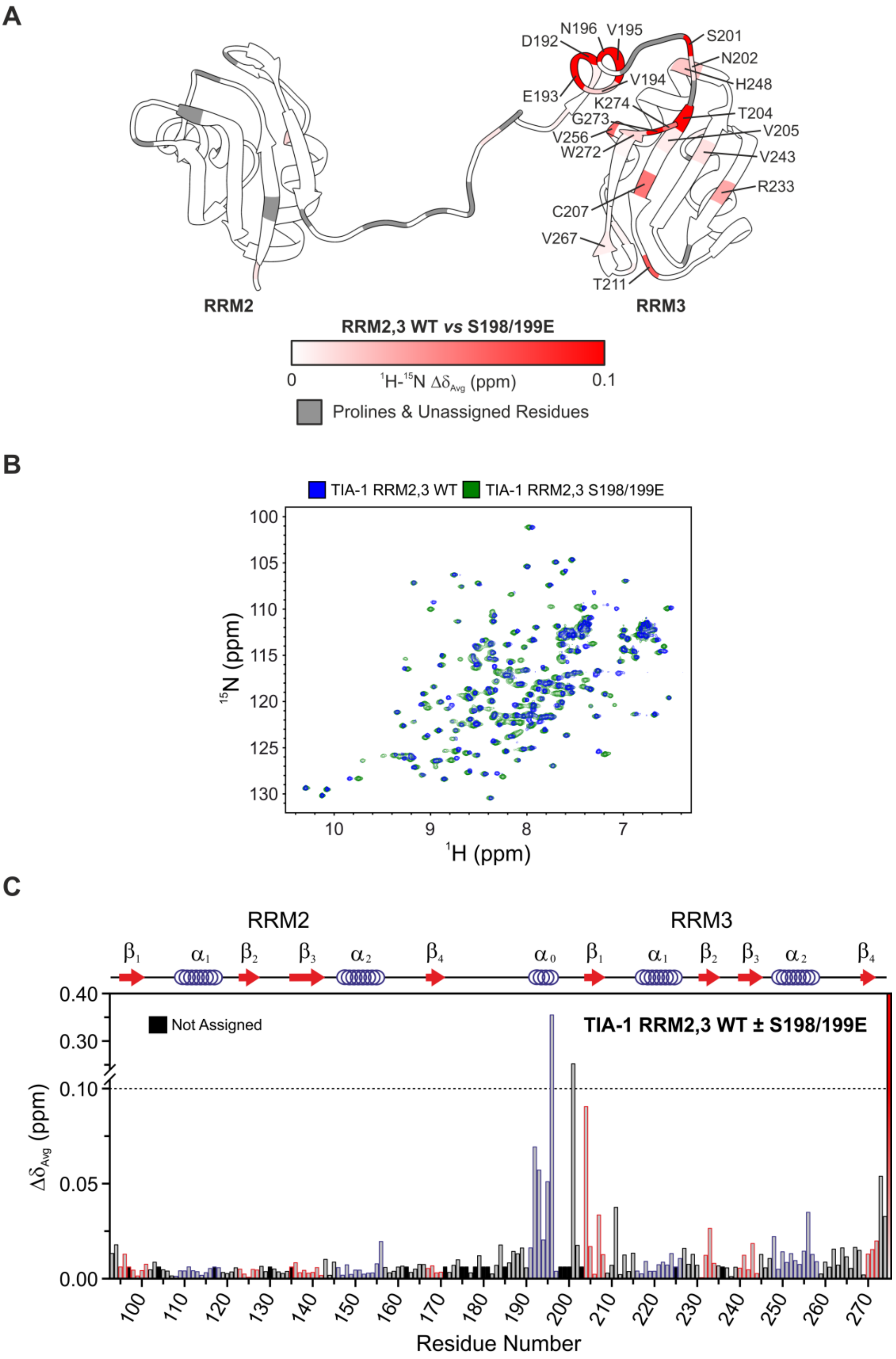


**Supplementary Figure S4. Effect of phosphomimetic double mutation S198/199E on TIA-1 RRM2,3 NMR signals. A)** Mapping of TIA-1 RRM2,3 residues (structure from ref. 29; PDB ID: 2MJN) perturbed by S198/199E mutations. Residues are colored according to their weighted-average chemical-shift perturbations ($\Delta\delta_{Avg}$) using a red gradient from 0 to 0.1 ppm. Prolines and not assigned residues are indicated in grey. **B)** $^{1}$H-$^{15}$N HSQC superimposed spectrum of TIA-1

RRM2,3 WT (blue) and TIA-1 RRM2,3 50PRD S198/199E (green). **C)** Bar graph of weighted-average chemical-shift perturbations ($\Delta\delta_{Avg}$) for $^{15}N$-labeled TIA-1 RRM2,3 due to S198/199E mutations, plotted versus residue number. Prolines and not assigned residues are indicated with black bars. Secondary structural elements are represented on top of each graph: blue overlapping circles for α-helices, red arrows for β-sheets and black flat lines for unstructured regions. In addition, the outline of the bars is coloured according to the secondary structure of the residue to which it belongs.

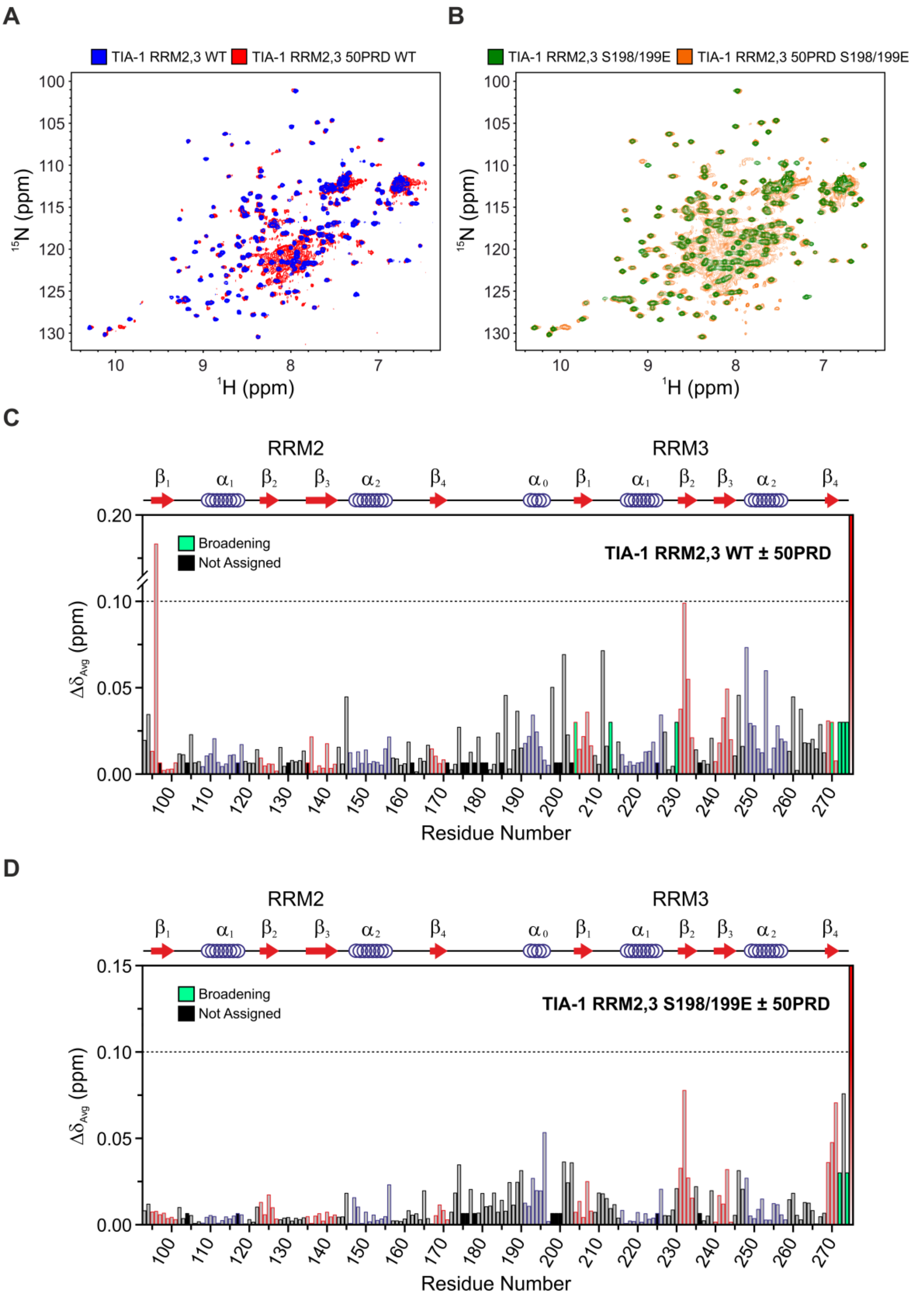


**Supplementary Figure S5. Effect of the first half of the PRD on TIA-1 RRM2,3 NMR signals. A-B)** $^{1}$H-$^{15}$N HSQC superimposed spectrum of TIA-1 RRM2,3 (blue or green) and TIA-1 RRM2,3 50PRD (red or orange), for WT (**A**) and S198/199E (**B**) constructs. **C-D)** Bar graphs of weighted-average chemical-shift perturbations ($\Delta\delta_{Avg}$) for $^{15}$N-labeled TIA-1 RRM2,3 WT (**C**) or S198/199E (**D**), due to the incorporation of 50PRD, plotted versus residue number. Residues with signals broadened beyond detection are indicated with green bars, whereas prolines and not assigned residues are indicated with black bars. Secondary structural elements are represented on top of each graph: blue overlapping circles for α-helices, red arrows for β-sheets and black flat lines for unstructured regions. In addition, the outline of the bars is coloured according to the secondary structure of the residue to which it belongs.

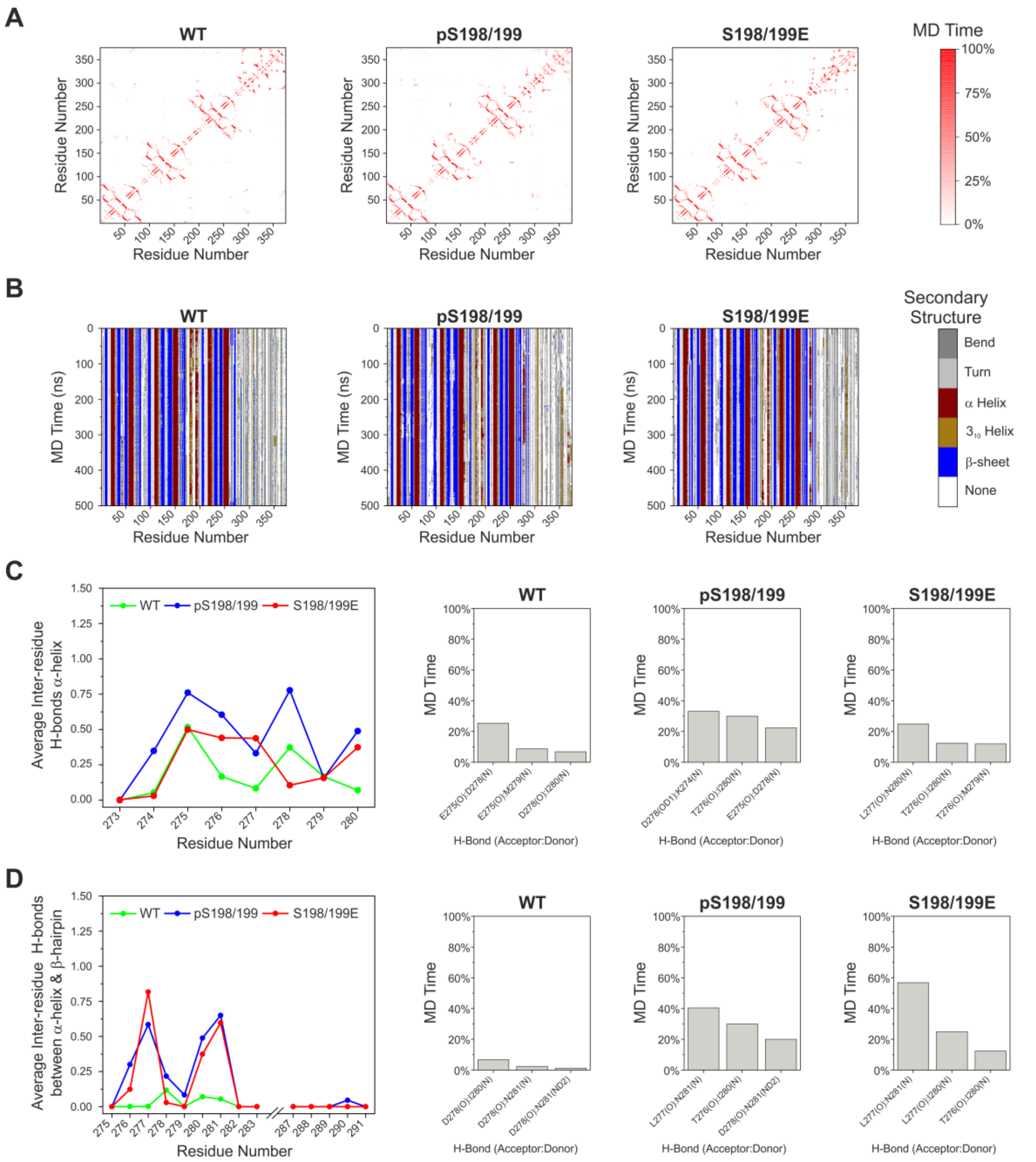


**Supplementary Figure S6. MD analysis of intramolecular contacts, secondary structure and hydrogen bonding for TIA-1 WT, pS198/199 and S198/199E models. A)** Inter-residue contact maps obtained from the MD simulations of TIA-1 and covering the whole protein. A red gradient has been used to indicate the percentage of MD time in which a given contact occurs (cut-off of 6 Å). Only contacts between residues separated by at least 3 residues in sequence have been computed. **B)** Time evolution of the secondary structure of TIA-1 during MD simulations, also covering the whole protein. **C)** Comparison of the average number of hydrogen bonds between residues comprising or near the predicted α-helix (residues 273-280) preceding the β-hairpin identified in TIA-1, along MD trajectories (left panel). Additional bar graphs reveal the top 3 most frequent hydrogen bonds in each model. **D)** Comparison of the average number of hydrogen bonds between residues from the predicted α-helix (residues 275-279) and β-hairpin strands (residues 280-283, 287-291) at the beginning of the TIA-1 PRD, along MD trajectories (left panel). Additional bar graphs reveal the top 3 most frequent hydrogen bonds in each model.

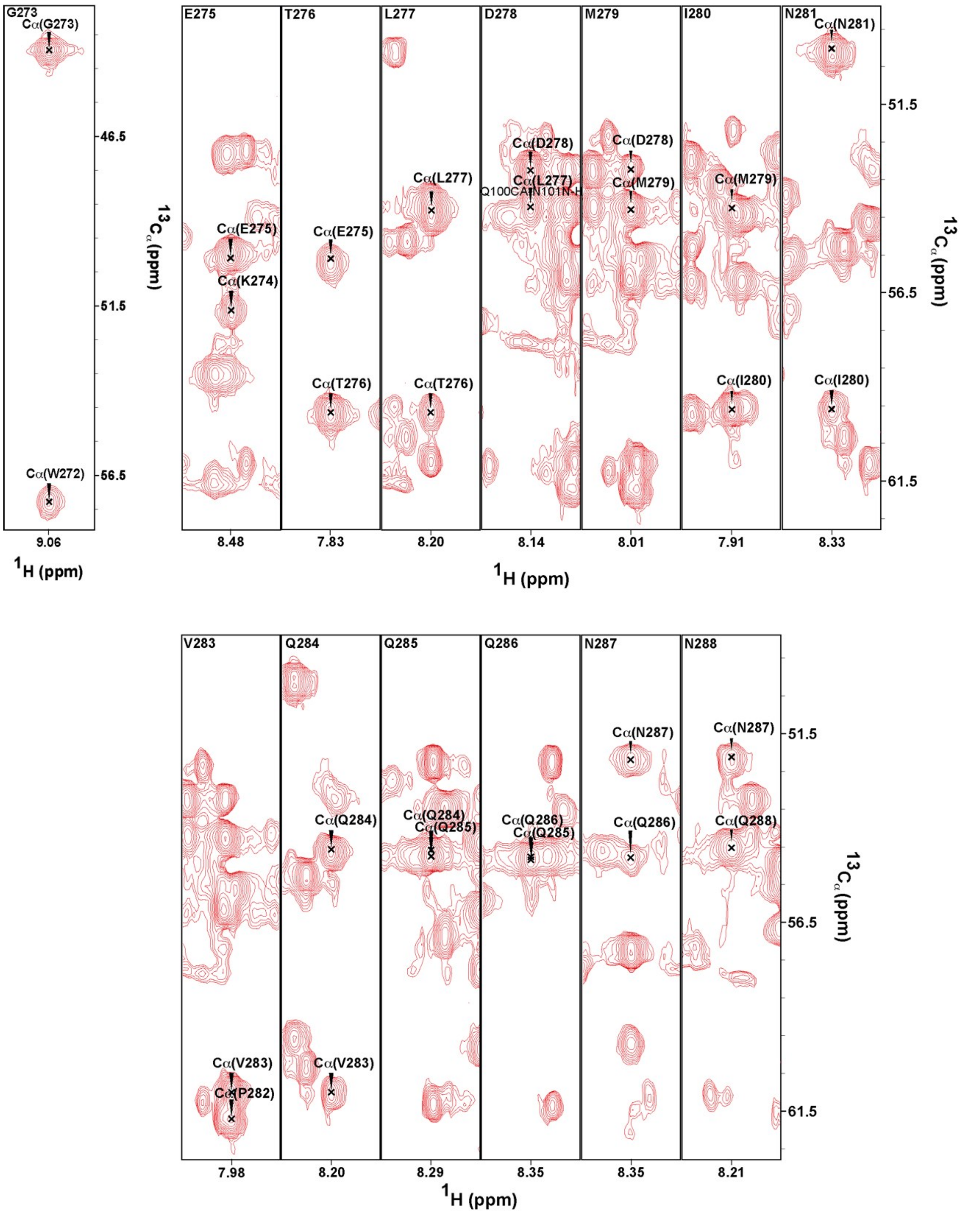


**Supplementary Figure S7. Strips of the 3D HNCA spectrum of TIA-1 RRM3 25PRD S198/199E illustrating the $^{13}C_\alpha$ chemical shifts for G273-Q288 residues located in the N-terminal quarter of PRD.** In each strip, the black cross indicates the chemical shift of the $^{13}C_\alpha$ of residue “n” or “n-1”.

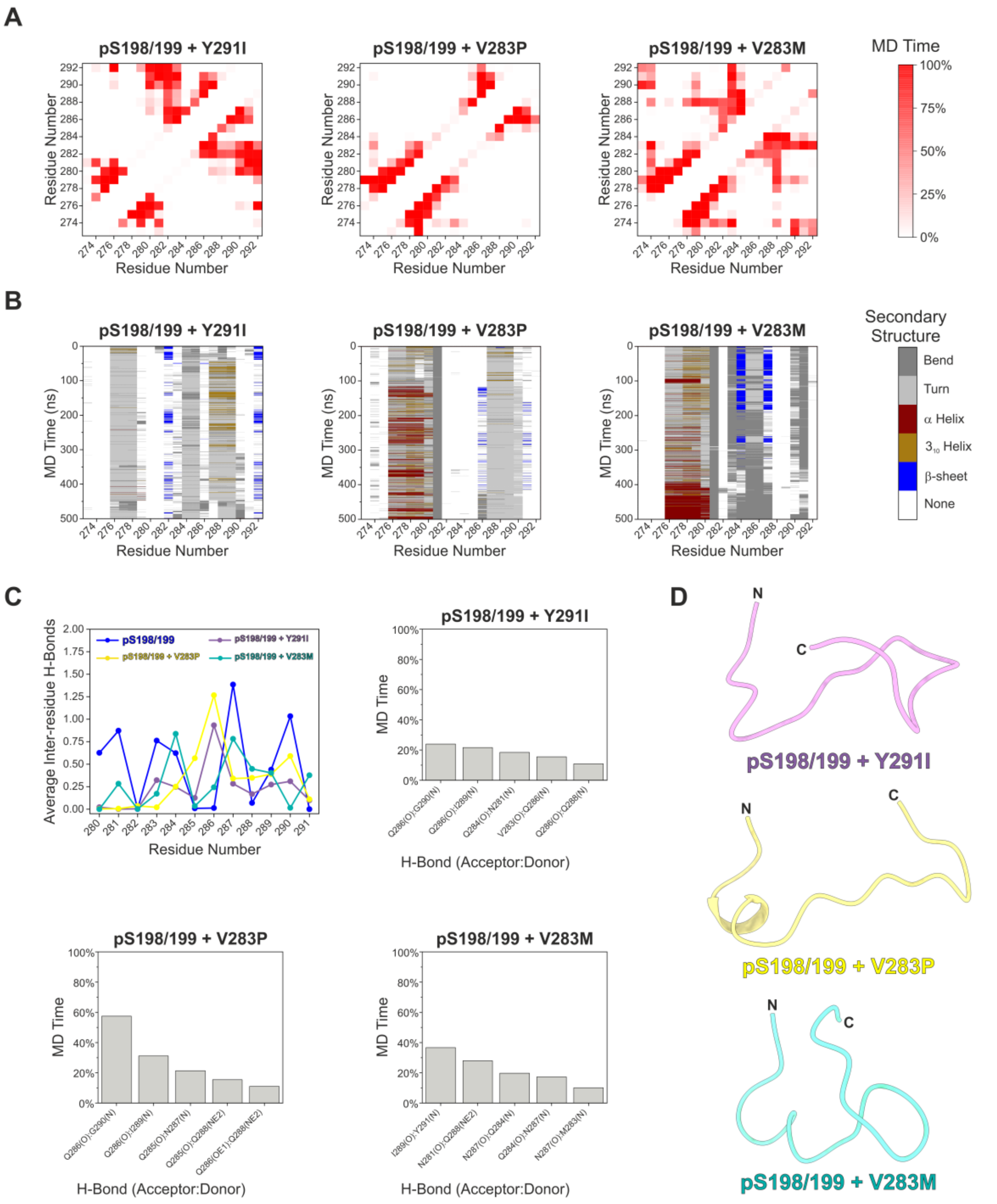


**Supplementary Figure S8. MD analysis of intramolecular contacts, secondary structure and hydrogen bonding for S198/199-phosphorylated TIA-1 Y291I, V283P and V283M models. A)** Inter-residue contact maps obtained from the MD simulations of TIA-1 and covering the first 20 residues of the PRD. A red gradient has been used to indicate the percentage of MD time in which a given contact occurs (cut-off of 6 Å). Only contacts between residues separated by at least 3 residues in sequence have been computed. **B)** Time evolution of the secondary structure of TIA-1 during MD simulations, also centered on the first 20 residues of the PRD. **C)** Comparison of the average number of hydrogen bonds between residues of the predicted β-hairpin motif (280-291) of TIA-1 along MD trajectories (top left panel). Additional bar graphs reveal the top 5 most frequent hydrogen bonds in each model. **D)** Representative ribbon structures of the most populated clusters from the MD simulations of TIA-1, showing the β-hairpin region. Clustering of trajectory frames was based on RMSD (residues 191-297).

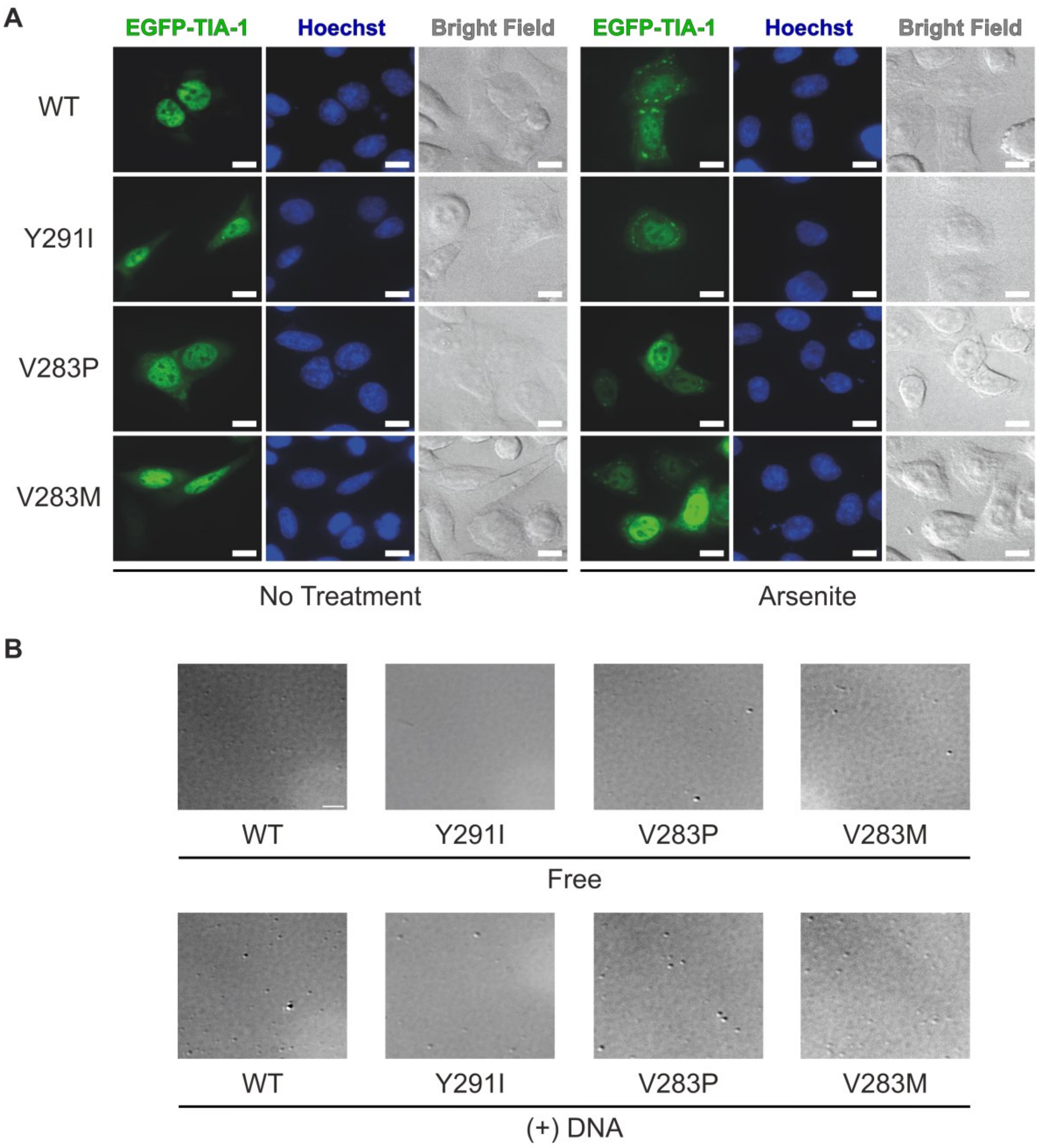


**Supplementary Figure S9. Representative microscopy images from experiments with TIA-1 β-hairpin mutants. A)** Representative fluorescence microscopy images of HeLa cells transfected with different EGFP-TIA-1 constructs (green fluorescence) under homeostatic and oxidative stress conditions. Nuclei were stained with Hoechst (blue fluorescence). Scale bars are 10 µm. **B)** Representative bright-field microscopy images of different TIA-1 full-length constructs used in droplet formation assays, in the presence and absence of the TC5 ssDNA oligo. Scale bar is 5 µm.

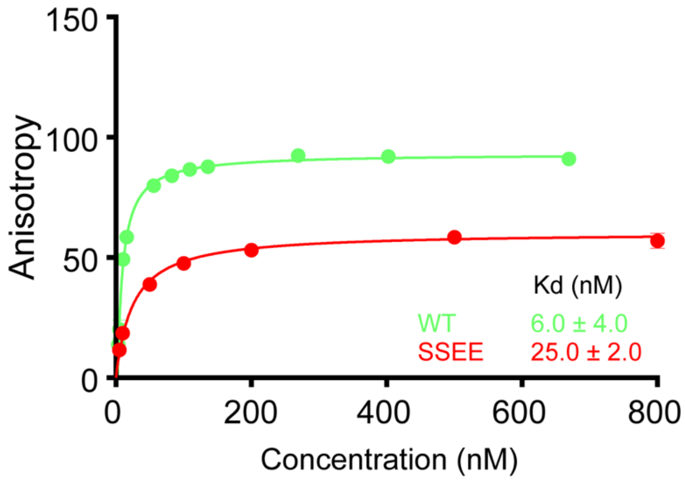


**Supplementary Figure S10. Fluorescence anisotropy binding curves for TIA-1 RRM2,3 WT and S198/199E**. Binding curves for TIA-1 constructs *versus* UC1 RNA. Data points are representing triplicate samples and the errors shown are standard errors of the mean (SEM). The binding data were fit by the 1:1 Langmuir binding model to derive the $K_D$.

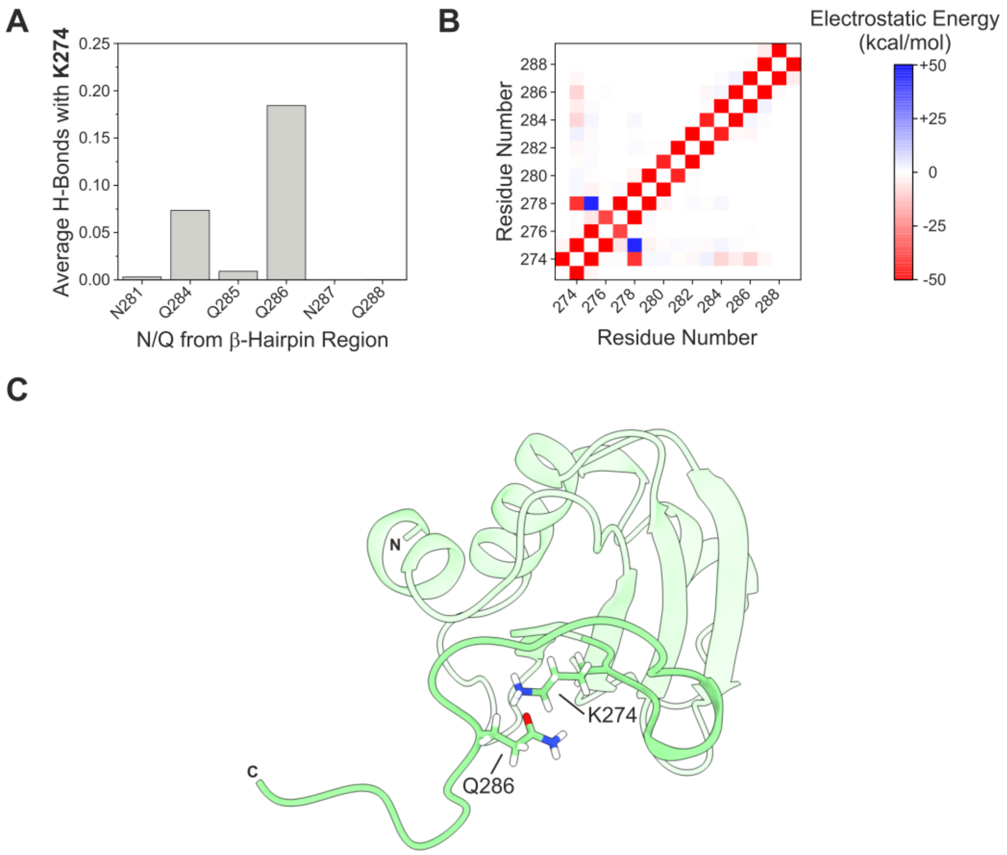


**Supplementary Figure S11. Hydrogen bonding network and electrostatic interactions of TIA-1 residues potentially involved in β-hairpin formation. A-B)** Bar graphs showing the average number of hydrogen bonds (**A**) and heatmap showing the pairwise average electrostatic interaction energy (**B**) between K274 and N/Q residues from the β-hairpin identified in TIA-1, along the MD simulation of the WT protein. The nature and magnitude of the electrostatic forces are represented by a diverging color palette, where attractive interactions appear in red and repulsive interactions in blue. **C)** Representative ribbon structure of the most populated cluster from the MD simulation of TIA-1 WT, showing the electrostatic interaction between K274 and Q286. Clustering of trajectory frames was based on RMSD (residues 191-297).

#### ImageJ macros

'Nuclei Mask'

```
run("Grays");
run("Invert");
setAutoThreshold("Default dark");
//run("Threshold...");
setAutoThreshold("Huang dark");
//setThreshold(233, 255);
setOption("BlackBackground", false);
run("Convert to Mask");
run("Close");
run("Convert to Mask");
run("Analyze Particles...", "size=50-500 display exclude clear add");
```

'SGs Analysis'

```
roiManager("Select", 0);
run("Duplicate...", "duplicate channels=2");
run("Enhance Contrast", "saturated=0.25");
run("8-bit");
run("Subtract Background...", "rolling=10");
run("Non-local Means Denoising", "sigma=15");
run("Auto Local Threshold...", "method=Bernsen radius=10 parameter_1=40 parameter_2=0 white");
run("Convert to Mask");
run("Watershed");
roiManager("Select", newArray(0,1));
roiManager("XOR");
roiManager("Add");
roiManager("Select", 2);
run("Analyze Particles...", "size=0.10-Infinity display exclude overlay");
```

Notes:

- ROI numbering: whole cell is 0, nucleus is 1 and cytoplasm is 2.
- The green fluorescence channel is number 2.
- The ImageJ Non-local Means plugin for denoising images must be downloaded (https://sites.imagej.net/Biomedgroup/plugins/) and placed in the ImageJ Plugin directory before using the macro.
- If ROIs disappear from the binary image after running the macro and the Results table does not appear, go to *Edit > Selection > Restore Selection*, and then run the *Analyze Particles* command.
- For optimal performance of the macro, it is highly recommended to carry out preliminary tests with several of the images to be analyzed in order to adequately adjust the parameters of each processing step. After finding the best settings, keep the macro unchanged throughout the analysis of all images to avoid any bias.

‘Droplets Analysis’

```
run("Duplicate...", "duplicate channels=2");
run("Enhance Contrast", "saturated=0.25");
run("Subtract Background...", "rolling=50 light");
run("Non-local Means Denoising", "sigma=15");
run("8-bit");
run("Auto Local Threshold", "method=Phansalkar radius=15 parameter_1=0 parameter_2=0");
run("Invert");
run("Remove Outliers...", "radius=5 threshold=25 which=Bright");
run("Convert to Mask");
run("Analyze Particles...", "size=0.10-Infinity circularity=0.30-1.00 add");
```

Buades, A., Coll, B. and Morel, J.-M. (2011). Non-Local Means Denoising. *Image Process. Line.* **1**: 208–212.
